# Mass Extinction Triggered the Early Radiations of Jawed Vertebrates and Relatives (Gnathostomes)

**DOI:** 10.1101/2025.09.11.673971

**Authors:** Wahei Hagiwara, Lauren Sallan

## Abstract

Most vertebrate lineages are first recorded from the mid-Paleozoic, well after their Cambrian origin and Ordovician invertebrate biodiversification events. This delay has been poorly understood and is usually attributed to sampling and long ghost lineages. We analyzed new databases of Paleozoic vertebrate occurrences, biogeography, and ecosystems, revealing that the Late Ordovician Mass Extinction (~444-443 million years ago) triggered parallel, endemic radiations of jawed and related jawless vertebrates (gnathostomes) in isolated refugia. Post-extinction ecosystems hosted the first definitive appearances of most major vertebrate lineages of the Paleozoic “Age of Fishes” (and today), following the loss of ubiquitous stem-cyclostome conodonts, nascent faunas of other gnathostomes, and pelagic invertebrates. Turnover and recovery patterns matched those following climatically similar events like the end-Devonian mass extinction, including a post-extinction “gap” with low biodiversity. The prolonged 23 million year Silurian recovery, and the challenges of oceanic dispersal, likely further delayed the dominance of jawed gnathostomes for millions of years after the first fossil jaws.

## Introduction

The more than 75,000 living vertebrate species belong to just two clades, the many jawed gnathostomes (tetrapods and “fishes”) and the few jawless cyclostomes (lamprey and hagfishes)(*1, 2*). Stem-vertebrates first appeared more than 520 million years ago (Ma) in famous “Cambrian Explosion” faunas such as Chengjiang and Burgess Shale alongside the early members of other phyla (*2, 3*). Cyclostomes diversified shortly thereafter in the form of their ubiquitous marine stem-members, the conodonts (*3, 4*), in line with Cambrian and Ordovician invertebrate biodiversification events (*5, 6*). It is widely assumed that gnathostomes, particularly jawed forms, must have likewise diversified in the early Paleozoic (*7, 8*), based on mid-late Cambrian (521–487Ma; *3, 9*) estimated divergence dates for crown vertebrates (*10*) and scattered stem- and crown-gnathostome like fossils from the mid-late Ordovician (471–443Ma; *2, 7, 10, 11*). However, gnathostomes did not show up in any real abundance, or at all in most regions, until the Silurian (443–420Ma; *2, 8, 10, 12*). This implies a 50–100 million year gap in the record of most lineages, particularly among jawed fishes. The gap is usually attributed to poor sampling or environmental constraints (*3, 7, 8, 12*), ignoring the global abundance of preservationally-similar conodonts and ecologically-analogous mobile, pelagic invertebrates (*2, 10*).

The early Paleozoic “gnathostome gap” is punctuated by global events that are significant in the evolution of marine biodiversity, but have received only passing consideration by vertebrate paleontologists (*2, 8, 10, 12*). This includes the double-pulsed Late Ordovician Mass Extinction (LOME, 445–443 Ma), a “Big Five” event marked by prolonged global fluctuations in temperature, sudden polar glaciation, and sea level changes which drowned or marooned coastal faunas (*13–15*). The LOME is linked to significant losses in pelagic and predatory invertebrates (e.g. ammonoids and arthropods), as well as conodonts (*14*). In all these characteristics, the LOME is highly similar to another glaciation-linked event within a longer crisis interval, the end-Devonian mass extinction or Hangenberg Event (EDME; 359 Ma), which profoundly devastated vertebrate ecosystems (*2, 16, 17*). Following the LOME, the Silurian era (443–420Ma) comprises a 23–million year recovery interval, marked by reorganization and diversification of mobile invertebrate faunas (*15*), and coincident with the documented first appearance of most major gnathostome clades (*3,10*). The EDME was also followed by a similarly prolonged Mississippian (359–323Ma) recovery interval notable for the increased diversification and abundance of living gnathostome lineages (*2, 16, 17*). The parallels between the Late Ordovician and end-Devonian in terms of drivers and influence on mobile marine animals suggest a possible role for mass extinction in observed Ordovician-Silurian changes in gnathostome diversity.

A major stumbling block in reconstructing early vertebrate diversification has been insufficient and poorly-veted global occurrence databases (*2, 10*). To determine the relationship between the early Paleozoic gnathostome gap and the LOME, we compiled the most complete record of early to mid-Paleozoic gnathostome occurrences to date. Our database contained 1,156 occurrences for 449 gnathostome species stretching from the early Ordovician to the end-Silurian (487–420Ma), and 2,546 species records for the Devonian (420–359Ma; *18*). As a comparison, we downloaded conodont stem-cyclostome occurrences from the Paleobiology Database (*18*).

We used these to produce genus-level, stage-binned diversity curves for all vertebrates, 13 major gnathostome groups, and 5 major geographic regions over the Ordovician-Silurian (Figs. 1, 4), as well for jawed and jawless fishes over the Silurian-Devonian (Fig. 4). We also assembled curves for richness within each stage per million years to correct for variations in stage length (Fig. 1B). To account for sampling bias and observe regional trends, we sorted gnathostome species occurrences into 168 Ordovician-Silurian assemblages (faunas) to detect changes in local species richness within major clades and overall faunal composition across time and environments (Fig. 2; *16*). Finally, we tracked gnathostome biodiversity across 5 major regions to determine trends in biogeography (Fig. 3).

**Figure 1.**
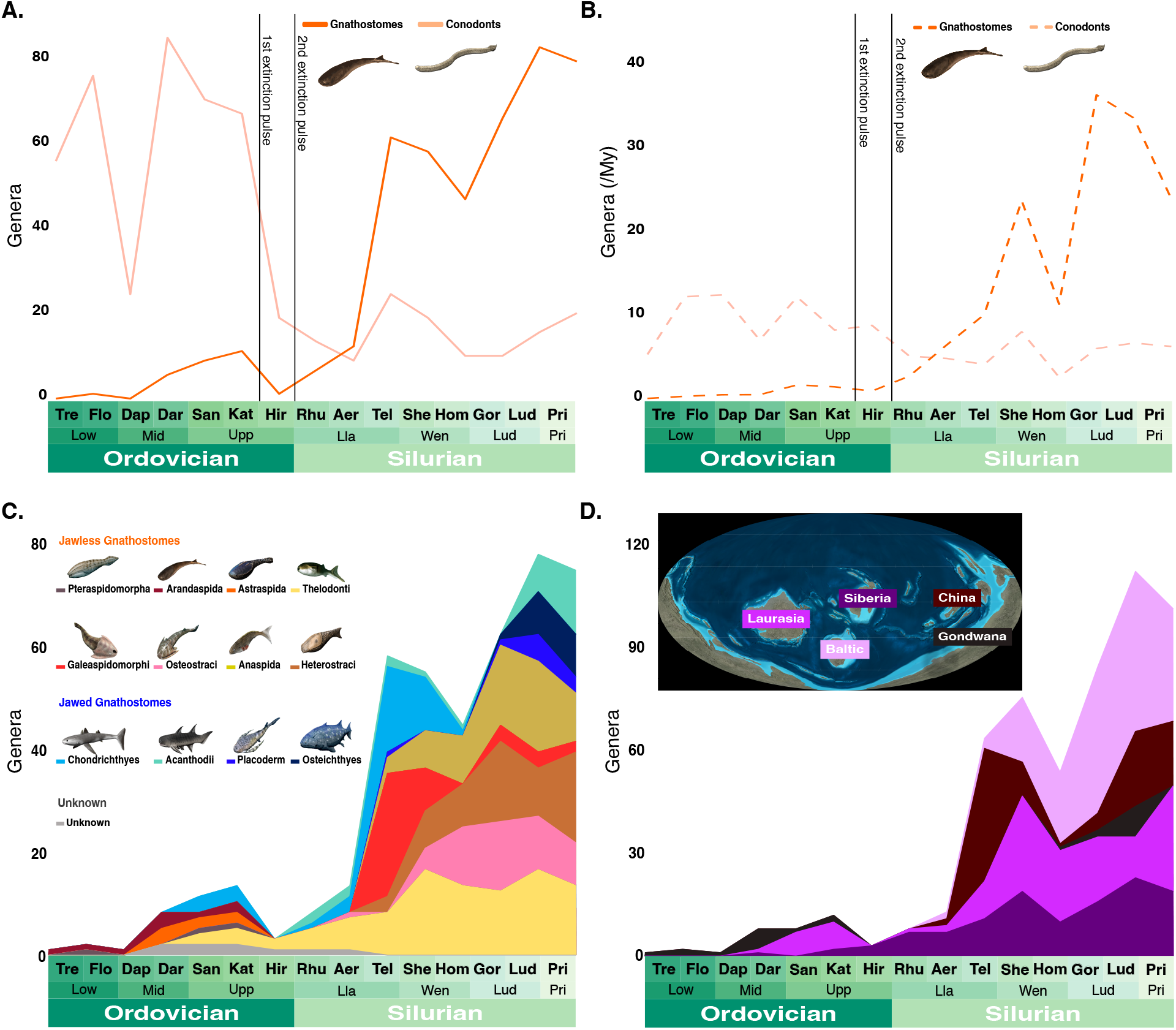
Diversity curves for gnathostomes from the Ordovician to Silurian. **(A)** Genus-level, stage-binned curves of global taxonomic richness for gnathostomes (N = 417) and conodonts (N = 480). Timescale based on *9*, extinction lines based on *14*. **(B)** Diversity curves based on richness per million years in each stage. **(C)** Diversity curves for genus-level richness per stage in 13 Paleozoic gnathostome classes (Fig. S1). **(D)** Diversity curves for stage-binned gnathostome genus-level richness in 5 major regions (Fig. S1). Map used with permission ©Deep Time Maps. Vertebrate reconstructions by Nobu Tamura.

**Figure 2.**
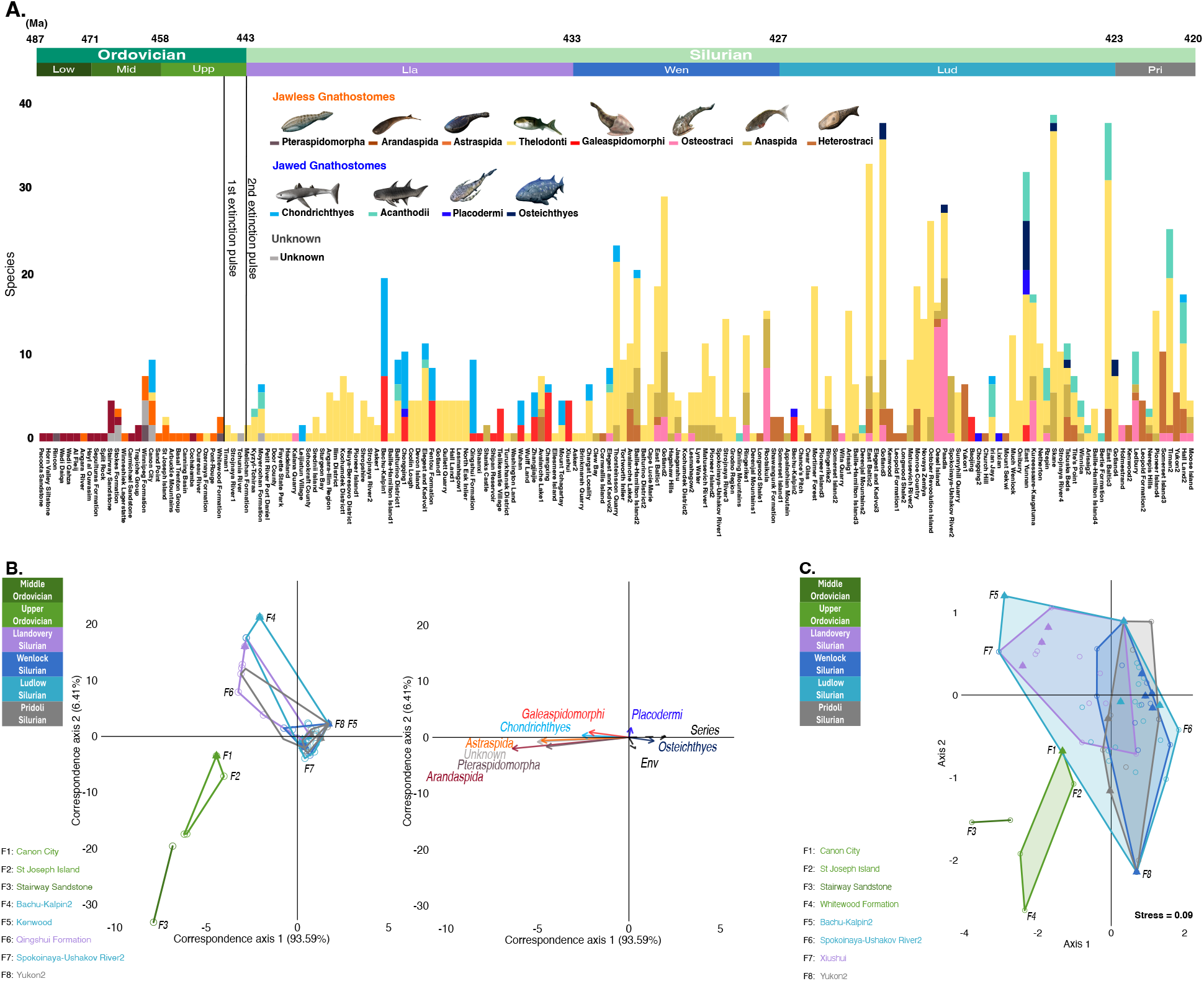
Change in the composition of gnathostome faunas through time. **(A)** Histogram of species-level faunal composition for all 168 Ordovician-Silurian assemblages (1,156 species-level occurrences) (Dataset S1). Species are binned into 13 major gnathostomes groups as in Fig. 1. The geological timescale (*9)* corresponds to the series-level age of each site, as some assemblages span multiple stages. Extinction pulses based on *18*. **(B)** Canonical Correspondence Analysis (CCA) of group-binned species diversity for a subsample of 101 localities containing at least 3 species (1,076 species occurrences). Filled triangles indicate marine sites, while open circles indicate non-marine. **Left:** Assemblage ordination plot (N = 101), with the most disparate Ordovician and Silurian faunas on each axis indicated (N = 9). **Right:** Group ordination plot (N = 13). Biplot arrows represent the 8 gnathostome groups with the highest contributions to assemblage position, and two explanatory variables (series or geological time; environment) based on their correlation with taxon distribution. **(C)** Non-Parametric Multidimensional Scaling (NMDS) based on gnathostome group species richness at assemblages and Bray–Curtis distances, with the 8 most disparate Ordovician and Silurian assemblages on each axis indicated. Alternative grouping combining Chondrichthyes and Acanthodii shows the same result (*18)*. Vertebrate reconstructions by Nobu Tamura.

**Figure 3.**
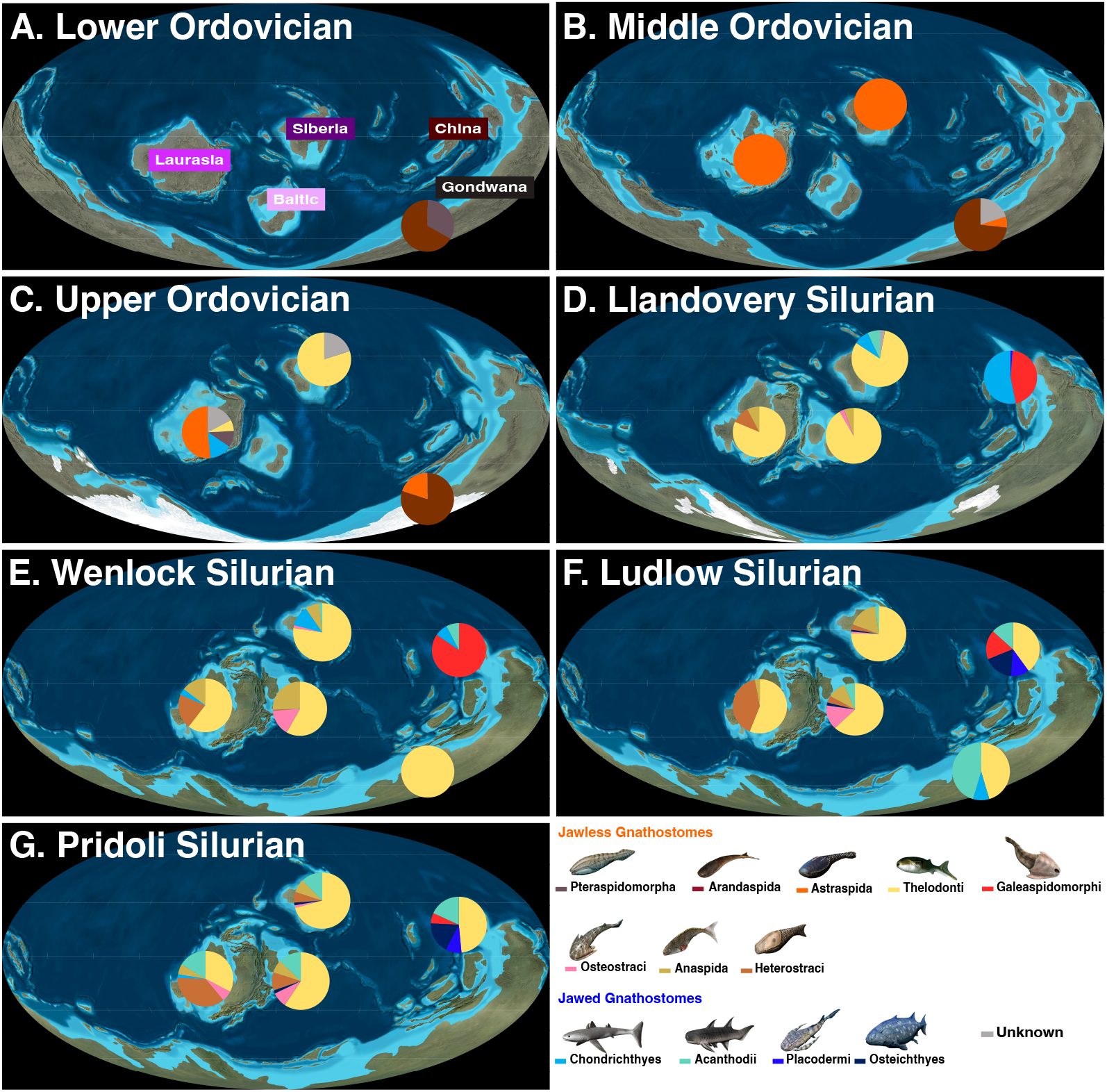
Genus-level richness by geological series for 13 gnathostome groups in five Paleozoic geographic regions in (Gondwana, Siberian, Laurasia, Baltic, China). **(A)** Lower Ordovician (487 Ma-471 Ma), (B) Middle Ordovician (471 Ma-458 Ma), (C) Upper Ordovician (458 Ma-443 Ma), (D) Llandovery Silurian (443 Ma-433 Ma), (E) Wenlock Silurian (433 Ma-427 Ma), (F) Ludlow Silurian (427 Ma-423 Ma), (G) Pridoli Silurian (423 Ma-420 Ma). Maps used with permission ©Deep Time Maps. Vertebrate reconstructions by Nobu Tamura.

**Figure 4.**
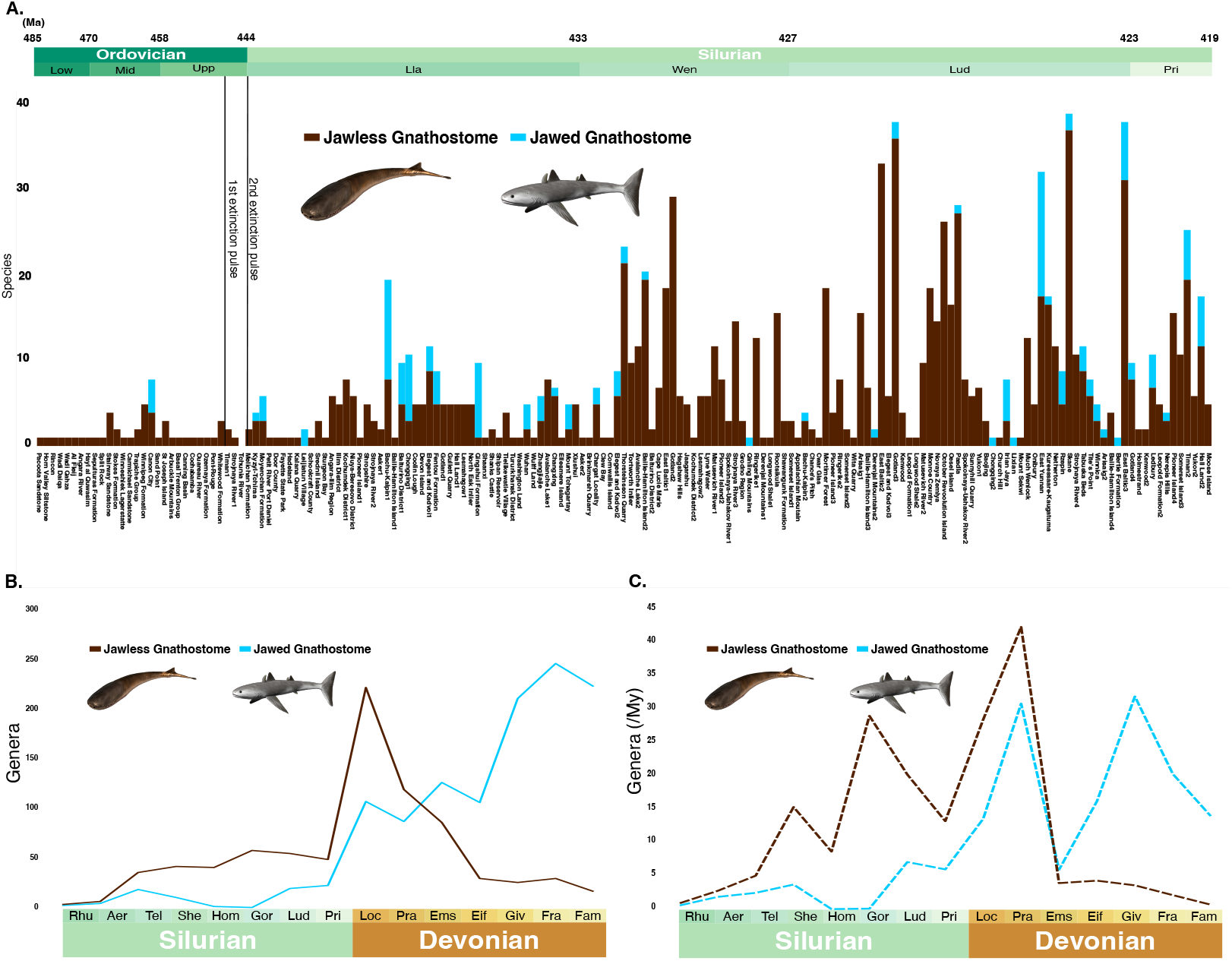
Comparison of jawed and jawless gnathostome diversity trends over the mid-Paleozoic. **(A)** Histogram showing species level richness for jawed and jawless gnathostomes for assemblages from Ordovician to Silurian, using same dataset as in Fig. 1A excluding “unknown” species (N = 1,143 species occurrences). Group assignment based on taxonomic attribution, rather than direct evidence for jaws. **(B)** Global diversity curves for stage-binned genus richness for jawless (N = 830) and jawed (N = 1,188) gnathostome groups for the Silurian and Devonian, with timescale based on *9*. **(C)** Global diversity curves for stage-binned genus richness per million years. Vertebrate reconstructions by Nobu Tamura.

## Results

First, we found that gnathostomes and cyclostome (conodont) genera exhibited a classic double wedge pattern with turnover centered on the two end-Ordovician extinction pulses (Fig. 1A). Our stage-binned diversity curves suggested high conodont richness in the Ordovician and extreme losses over the end-Katian (445Ma; Fig. 1A). However, time correction blunted some of the LOME impact, suggesting stability and then a muted decline over the end-Hirnantian (443Ma) that settled at a slightly lower average in the Silurian (Fig. 1B). This is in line with prior reports of conodont turnover at consistent richness levels during the LOME interal (*14*).

Conodont genus richness never recovered to Ordovician peaks even in longer intervals, but settled below a maximum of 20 genera per stage, a limit which held until their extinction in the Triassic (*6*). In contrast to conodonts, gnathostome diversity was relatively low throughout the Ordovician, and even lower in time-corrected curves, with a slight increase in the Darwillian to Katian (469–445 Ma; Fig. 1A and 1B). Gnathostomes suffered apparent losses over the Katian, although this may be also have been a temporal artifact (Figs. 1A, B). We found that the entire Silurian contained a prolonged 23 million post-extinction diversification (or recovery) interval for gnathostomes, starting with 3-5 million years of very low global richness (Figs. 1, 4). Gnathostome global richness reached ever higher peaks (Figs. 1, 4B, C), and assemblages demonstrated greater richness and global homogeneity towards the end of the recovery interval (Fig. 2).

To determine whether the global diversification of Silurian gnathostomes results from LOME-related causes, we examined our records for Ordovician-Silurian clades, faunas and regions in more detail (*18*). Based on our faunal compilations (assemblages), the apparent increase in Late/Upper Ordovician gnathostome diversity (Fig. 2A) was mostly driven by two well-sampled intertidal faunas in North America (Canon City and Winnipeg)(Figs. 2A, 3C), one of which produces scales sometimes attributed to the earliest known jawed chondrichthyans (Fig. 2A)(*2, 3, 7, 10*). There was also an increased number of low richness assemblages containing members of Ordovician-specific jawless clades with clear habitat restrictions (*10*). Arandaspids (e.g. *Sacabambaspis*) only occurred in intertidal and subtidal areas along Gondwanan coastlines (Fig. 3)(*10, 18*). In contrast, astraspid pieces have been the dominant gnathostome fossils from middle Ordovician sites at then-equatorial shallow marine areas of Laurasia and Siberia, joined by thelodont scales in assemblages from near the end of the period (Fig. 3) (*19*). The biogeographic division between Northern and Southern continents was reflective of similar separation in invertebrate faunas, linked to intervening deep seas and unfavorable, East-West currents (*20*). Within these areas, the few well-sampled Ordovician gnathostome assemblages were highly homogeneous and distinct compositionally from post-extinction ecosystems in our ordinations, with the possible exception of the Chondrichthyes (“shark”) scale-bearing Canon City assemblage in our NMDS plot (Fig. 2D) (*18*).

Any nascent Ordovician gnathostome diversification appears to have been disrupted by the first glaciation pulse at the end of the Katian (Figs. 1C, 2A). Prior work has found that intrapulse Hirnantian mobile invertebrate and conodont faunas were a diverse mix of Ordovician relicts and diversifying species (*14*). We recovered only three Hirnantian gnathostome assemblages from Siberia, consisting of isolated thelodont scales belonging to just one or two form taxa as well as ichthyoliths unattributable to any prior species (Figs. 2A, 3C). The initial glaciation pulse marked the latest appearance of all more robust and armored Ordovician gnathostomes, including astraspids, arandaspids, and “Pteraspidimorpha” remains which may belong to their relatives (Figs. 3A-C). Almost all of Gondwana has been bereft of gnathostomes records from the entire Hirnantian and subsequent 10–15 million years of the Silurian, after which richness remained low (Figs. 2A, 3D) (*21*). This is observed even in areas which feature an otherwise robust Paleozoic record and have been subject to intensive sampling, such as Australia (Figs. 1D, 3) (*18, 21*). The gnathostome-depleted region was similar to the maximum extent of glaciation in the LOME (*14*). While some prior authors have proposed an “Out of Gondwana” circumpolar hypothesis for Silurian gnathostome diversification (*20*), this was ruled out by apparent extinction-related extirpation.

Following the LOME, gnathostomes exhibited fundamentally changed faunal and biogeographic diversity suggestive of deep losses. The earliest Silurian age, the ~3 million year Rhuddinian era (443–440.5Ma) of the Llandovery epoch, is coincident with “Talimaa’s Gap”, a term used for a previously inferred but neglected interval of low or missing vertebrate diversity (particularly in Gondwana as above; *10, 12, 21, 22*). We confirmed this quantitatively in our dataset, where Rhuddinian diversity increased only slightly from the Hirnantian (Fig. 1). The few Rhudinnian and early Aeronian (440.5–438.5Ma) age gnathostome-bearing sites produced almost only ichthyolith form taxa (Fig. 2A). Tropical Laurasian and Baltic regions were dominated by thelodonts in line with the nascent assemblages of the Hirnantian (Fig. 2A, 3D). In Siberian localities, thelodonts were joined by novel scales assigned to “chondrichthyan” or “acanthodian” form taxa such as tsunacanthids and elegestolepids, which differed from Ordovician types and disappeared after the Llandovery (Figs. 2A, 3D) (*23*).

All other major gnathostome groups made their first Silurian appearances in the Aeronian or Telychian (438.5–433Ma), 3–10 million years after the LOME. This occurred in line with increases in global genus richness (Figs. 1A, 1B). Most jawless stem-gnathostome clades, including osteostracans, galeaspids, and heterostracans (excluding Ordovician “pteraspidimorpha”) enter the Silurian record from separate regions (Fig. 3D). These groups exhibited initally low levels of richness and abundance ahead of more consistent diversification in the later Silurian (Fig. 1D). For example, while the earliest known osteostracan headshield came from an Aeronian quarry in Baltic Estonia, there has been a subsequent gap until the Wenlock (433–427Ma) of the same region (Figs. 2A, 3D, 3E). The Silurian emergence of heterostracans was even further delayed, as these initially appeared in the late Telychian record of Arctic Canadian Laurasia. This created a 10–million year gap (or ghost lineage) from last pre-extinction occurrence of similar, potentially-related “Pteraspidimorpha” and astraspids in Katian of North America (Figs. 2A). Galeaspids made their first appearance in the Aeronian of China and remained in this isolated region until their extinction in the Devonian (Figs. 1C, 2, 3D-F).

Chinese galeaspid ecosystems of the Aeronian and early Telychian also contained the earliest definitive evidence for jawed gnathostome lineages (*8, 24*). In our ordinations, chondrichthyan-galeaspid assemblages were distinct from Ordovician localities and early Silurian Laurasian and Baltic faunas, as well as a thick cluster of late Silurian sites on axis 1 in our CCA and axis 2 in our NMDS (Figs. 2B-D). A distinct Chinese/Asian fauna persisted throughout most of the Silurian (Figs. 3D-F). Our results suggest that the initial diversification of jawed fishes, and the origins of most major lineages, apparently occurred within a distinct refugium (*8*) in the first 10 million years post-extinction, even if stem-members first appeared elsewhere (*3, 7, 10, 11*). In the early Silurian, China was an equatorial offshoot of otherwise vertebrate poor Gondwana, separated from other continents by a deep sea (Figs. 3). While Ordovician China has not yet produced gnathostome material, it is possible that it hosted taxa found in nearby Australian coastal systems, such as arandaspids and the producers of jawed fish-like scale forms (*19, 21*). Survivors could have given rise to the local endemics in the post-extinction interval, after the extirpation of relatives from Australia itself (Figs. 1C, 3D). Endemic diversification of jawed fishes within Chinese ecosystems continued throughout the entire recovery interval (Figs. 1C,D, 2A, 3D-G, *8*). Ongoing geographic isolation in nearshore waters likely contributed to an observed relationship between Chinese assemblage size and the global richness of jawed fishes for most of the Silurian (Figs. 1C, 1D, 4B, 4C). However, divergence within Chinese faunas was not limited; heterogeneity and regional diversity increased as the recovery proceeded. Early “disaster” or “flux” ecosystems (*25*) of homogeneous, small stem-group chondrichthyans and galeaspids disappeared, replaced by a more diverse fauna of placoderms, osteichthyans and polyphyletic acanthodians (Figs. 1C, 3D-G). A Homerian gap between these faunas may represent either local sampling issues or a regional event, of which there were many in the unstable Silurian (Figs. 1C,D, 4B; *15*).

The spread of most gnathostomes from separate refugia occurred slowly during the Silurian recovery interval. Initially, dispersal over long distances was limited to specific lineages based on morphology (Fig. 2A, 3D-G; *10*). During the early-mid Silurian, faunas outside China were dominated by pelagic thelodonts (Fig. 2A and 3D) (*18*). Gnathostome communities were re-established in Gondwana by the Wenlock (Fig. 3E), but samples have consisted entirely of thelodont scales (Fig. 3E). Thelodonts also disrupted the distinctiveness of Chinese faunas through invasion by the Ludlow (427–423Ma; Fig. 3F), coincident with the disappearance of older scale- and spine-based jawed taxa (e.g. mongolepids) and a reduction in galeaspid richness mentioned above (Fig. 1C, 2A). Likewise, streamlined anaspids such as *Birkenia* spread to Wenlock Siberian and Baltic ecosystems within a short time after their initial appearances in the late Telychian of Laurasia (Fig 2A, 3D, 3E). In contrast, more environmental restricted, armored (macromeric) jawless lineages (e.g. galeaspids, osteostracans, heterostracans; *10*) remained limited to their natal continents for most of the Silurian, reaching adjacent lands only in the latest Silurian or Devonian in line with the formation of Pangaea (*15*), if at all (Fig. 3D-3G)

Most jawed fishes showed limited dispersal throughout the 23 million year recovery interval, remaining geographically restricted through the Pridoli (423–420Ma, Figs. 1C, 3D-G) like the coincident galeaspids and most armored jawless fishes elsewhere. Dispersal among even micromeric (small-scaled, flexible) jawed forms seems to have been delayed significantly relative to jawless fishes like thelodonts. Jawed fishes were apparently restricted to China and possibly Siberia throughout most of the Llandovery, with species in the latter consisting of a few isolated scales among a plethora of similar thelodont remains (Figs. 2A, 3E). By the Ludlow, Siberian ichthyoliths had been replaced by body fossils of acanthodians and the probable stem-Osteicthyan *Andreolepis*, which may have spread from China via prevailing currents (Fig. 3F). Acanthodians and other chondrichthyans continent moved rapidly to the Baltic following the counterclockwise conveyor belt of currents (*15, 20*). They also apparently moved against the flow via shallow seas to least one Gondwana Ludlow-age locality (New Guinea; Figs. 2A; 3F). Osteichthyans remained rare outside China throughout the later Silurian, with just one or two species present in scattered Baltic ecosystems (Figs. 2A, 3G). In contrast, acanthodian diversity increased dramatically in Pridoli ecosystems across all regions including Laurentia, perhaps enabled by the replacement of straits with shallows as Pangaea formed (Figs. 2G).

## Discussion

The Silurian reorganization and expansion of gnathostome biodiversity appears to have been driven by disruption to existing stable communities by the LOME, including extirpation of incumbents and resultant ecological release, in line with patterns across other mass extinctions (*16, 25, 26*). Conodonts and other pelagic victims of the LOME such as ammonoids and arthropods were the most likely candidates for such incumbents (Figs. 1A, B; *14*). However, the known ecomorphological diversity of Silurian gnathostomes far outstrips that inferred for pre-extinction conodonts based on just two Paleozoic lamprey-like body fossils recovered from after the initial extinction pulse (*27*). This could suggest that reorganization of vertebrate ecosystems after the LOME went beyond simple replacement (*10*). However, it is also possible Ordovician vertebrates exhibited higher ecomorphological diversity than realized, particularly given the almost total lack of body fossils and poor preservation potential for such lineages (*3, 7, 28*). The hagfish-like Hirnantian conodont *Promissum* was also poorly preserved (*27*), but it could represent a post-extinction body plan which was carried forward by later ecologically conservative crown-cyclostomes (*4*). In any case, Ordovician conodonts are known to have exhibited high levels of diversity in their feeding traits and lived in a range of environments (*10, 29*), so it is probable they filled a large set of ecological roles in Ordovician seas.

Ordovician gnathostomes might also have contained a greater diversity of lineages and ecomorphologies that ultimately succumbed to the LOME, rather than being simply early representatives of known Silurian taxa as previously assumed (*3*). North American and Australian Ordovician fossil sites have produced “chondrichthyan” and “gnathostome” scales of indeterminant lineage and rare histology for the Paleozoic (Fig. 2A, C; *3, 7, 11, 18*). These forms have been significant different from scales found in early Silurian assemblages in China and Siberia (*7, 23*), as well as those found on Silurian body fossils (*24*), despite being made of the same dermal hard tissues. A few Ordovician ichthyolith taxa were listed as “unknown” because they have combinations of traits that have led to attribution to different later lineages at different times (Data S1; *18)*, and so may represent stem-group experimentation (*3*). More completely known taxa from Ordovician “armored” gnathostomes, such as arandaspids and astraspids, exhibited a few traits shared with later lineages such as heterostracans, leading to their inclusion in clade-specific phylogenies (*3, 10*), but otherwise exhibited divergent forms and lack derived characters. There is space for undersampled, unknown diversity of gnathostomes within the widespread occurrence of broken pieces of “pteraspidimorphs” in coastal assemblages, but these remains have been unattributable because diverge from later taxa (Fig. 2A; *3, 10*). Even the few Ordovician thelodont scale-based taxa never occurred again after the second glacial pulse, replaced by more regular Silurian forms such as *Loganellia* (Data S1). Therefore, it is likely that there was a fully distinct Ordovician gnathostome fauna, even involving very early members of Silurian lineages and localized diversification in shallow waters (*10*). Yet, all known Ordovician taxa appear to have been eliminated or substantially changed over the LOME, suggesting complete extinction-related turnover within gnathostomes (Fig. 1A-C).

We found strong evidence for a prolonged post-extinction recovery (*26*), in which most Silurian gnathostome lineages diversified gradually during an initial period of otherwise very low diversity. Indeed, the Llandovery epoch (443–433Ma) featured delayed vertebrate appearances and reappearances in distinct, regionally homogenous and low diversity post-extinction faunas. This pattern is similar to vertebrate diversity during the post-EDME Tournasian (359–345 Ma; Fig. 2; *2, 16*). Given proximity to the end-Hirnantian glaciation event, Talimaa’s Gap is analogous to the “Romer’s Gap” diversity trough following the EDME glaciation and similar intervals after other events (*2, 16, 26*). With the exception of Siberia, many of early Llandovery faunas occurred in regions that have lacked good Ordovician gnathostome records such as China and the Baltic (Figs 1A, 1B). Likewise, early Silurian rocks near productive Ordovician sites in Gondwana and elsewhere have not yet produced gnathostome samples (Fig. 3A-D). This suggests extreme biogeographic shifts throughout the Hirnantian and early recovery interval. In line with observations after other mass extinctions (*16, 27*), there was very low initial diversity and richness in most places during the first 3–5 million years the recovery (“Talimaa’s gap”, as above; *10*), and even the rest of the 10 million year Llandovery in some areas. Any surviving populations likely fell below the critical mass needed for preservation (*27*). Such low local diversity is supported by the fact that most known early Llandovery “assemblages” consist of less than 5 species in only 1 or 2 groups.

We observed a high level of endemism in gnathostomes from the very beginning of the Silurian, a stark change from the wide geographic ranges observed for Ordovician vertebrates. Regional distinctions in faunal composition (Fig. 1D, 3) suggested that ecological reorganization and diversification initially occurred within specific, long-lasting extinction refugia (*30*). This allowed the parallel emergence of novel species in different faunas, and perhaps some degree of early experimentation in form. For example, many Rhudinnian ecosystems contained distinct taxa which do not appear either earlier or later on, such as mongolepid chondrichthyans and specific thelodont genera (Fig. 2A; *7, 18*). These may be interpreted as short-lived “disaster taxa,” or residents of transient “disaster faunas” or “flux ecosystems,” both common post-extinction phenomena in invertebrate records (*27, 28, 31*). We also found that each early Silurian region featured a distinct set of novel benthic forms, such as galeaspids and osteostracans. These exhibited divergence within a highly-restricted nearshore environment (*10*), alongside novel pelagic gnathostomes such as acanthodians and anaspids. Thus, while gnathostome diversity remained low for millions of years after the LOME, separate refugia enabled high levels of disparification and divergence in protected endemic areas. The post-extinction ecosystems created the template for widespread gnathostome communities in the Devonian “Age of Fishes” and in modern oceans.

## Materials and Methods

### Dataset assembly procedure

A dataset of gnathostome occurrences from the Ordovician-Silurian interval was compiled from literature. The details of sources are provided in Data. S1, S3, S4, specifying which were used to determine taxonomic classification, age, discovery location and environment of their habitat. Taxonomic information was primarily based on the latest data available in each paper, as well as general references such as Sepkoski’s compendium (*6*), *Family-Group Names of Fossil Fishes* (*32*) and the *Fossiilid*.*info* database (*33*). The dataset includes 1,156 occurrences, 449 species-level records and 219 genus-level records from the Ordovician–Silurian interval. For genus-level analyses, indeterminate records (gen. indet.) were excluded, while they were retained for analyses of faunal composition (see below). Fossil occurrences assigned to 168 distinct faunal assemblages on the basis of shared localities, geological settings, environment, and age, following the procedure of Sallan and Coates, 2010 (*16*, Data. S1). Similarly, taxonomic and age information for Devonian occurrences were compiled using the same approach (Data. S5); however, faunal assignments of Devonian were not included in this study. Additionally, the occurrences of conodonts across each interval were downloaded from the Paleontology Database and compiled by stage (PBDB) (*34*) (Data. S6–7) resulting in a total of 480 genus-level records for the Ordovician-Silurian (Figs. 1A, 1B). This was augmented by genus-level records from Sepkoski’s compendium (*6*) in order to fit Silurian diversity trends in the context of the later record for the clade until their end-Triassic extinction (*2*).

### Taxonomic group assignments for occurrences

Each gnathostome record in each assemblage was classified into 1 of 12 categories representing major gnathostome lineages or groups (*10;* Data S1), based on their assignments in the literature including 8 Paleozoic jawless fish classes (Arandaspida, “Pteraspidomorpha”, Astraspida, Thelodonti, Galeaspidomorphi (Galeaspida), Osteostraci, Heterostraci, and Anaspida) and four Paleozoic jawed fish classes (“Chondrichthyes”, “Acanthodii, “Placodermi,” and Osteichthyes). Additionally, an “Unknown” category was assigned to taxa whose exact classification remains undetermined or controversial in the literature, but which are distinct from co-occurring species and have valid genus names. Of the other categories, two may be paraphyletic or polyphyletic (Pteraspidimorpha and Acanthodii) but represent distinct sets of similar species and have been used as historical categories. We explain the usage and assignment of “Unknown”, “Pteraspidimorpha”, “Acanthodii”, “Chondrichthyes,” and “Placodermi” in further detail below.

The “Unknown” category in this study was primarily used for enigmatic microfossil records from the Ordovician and Silurian. This category included *Skiichthys halsteadi* and *Eleochera glossa* from our “Winnipeg Formation” and “Canon City” assemblages in the United States (Data S1) (*35, 36*). The former species has been suggested to have affinities with Acanthodii or Placodermi (*8*) but its classification remains uncertain (*37*). The latter species exhibits affinities with Chondrichthyes or Anaspida; however, its precise taxonomic placement has not been determined (*38*). Additionally, Ordovician taxa identified from microfossil records and previously attributed to “Chondrichthyes”, such as those in the “Stairway Sandstone”, “Stokes Formation”, and “Winnipeg Formation” assemblages, were also re-assigned to the “Unknown” category. This was due to uncertainty expressed by the same authors that made these assignments in later papers, as well as well as doubts expressed by other workers (*3, 10*, Data S1). The exceptions are taxa from “Canon City”, which have been classified as stem-Chondrichthyes continuously in the literature and by the same authors, despite some minor uncertainty (*3, 10, 38;* see discussion of “Chondrichthyes” below). *Tesakoviaspis concentrica*, a scale taxon recovered from “Tchunia River” and “Moyerochan Formation” in Siberia during the latest Ordovician to earliest Silurian was also classified as “Unknown”. Although it has previously been assigned as Astraspida (*6*), histological evidence has raised doubts about this placement (*39*). *Dictyorhabdus priscus* from “Canon City” (*40, 41*) was excluded from the dataset, as it has not been confirmed to be a true vertebrate, despite suggestions of a weak affinity with other gnathostomes (*42, 43*).

“Pteraspidomorpha” was used for the Ordovician taxa *Pircanchaspis rinconensis* (*44*), *Pycnaspis splendens* and *Pycnaspis sp. cf. splendens* (*37*), as their placement in relation to Arandaspida and Astrapida remains unresolved, and they may represent distinct groups of Ordovician gnathostomes (see main text). Arandaspida, Astraspida, and Heterostraci are here treated separate groups because they have been regarded as subclasses within Pteraspidomorpha (*10, 32*). In addition, recent histological and phylogenetic work on arandaspids and astraspids has found that these were sister groups to the exclusively Silurian Heterostraci and further highlighted the distinctions between Arandaspida and Astraspida (*42, 45, 46*). Therefore, we decided to treat these subclasses as distinct groups in line with previous work (*10*).

“Acanthodii” were previously considered a distinct class and later a polyphyletic set of differentiated clades (e.g. Ischnacanthida, Acanthodida, Gyracanthida, Climatiida) with affinities to chondrichthyans, stem-gnathostomes, and osteichthyans (*2, 16*). The discovery of the maxillate placoderm *Entelognathus* changed early gnathostome phylogeny such that acanthodians became a set of stem-chondrichthyan clades with distinct sets of characters (*47*).

While a number of Silurian and Ordovician scale and spine-based taxa have been assigned as stem-”chondrichthyans” without clarification (*7, 11, 23*), some of the same authors have also recently assigned early Silurian body fossils as “acanthodian-grade” on the basis of a distinct body plan shared with the older classification and distinct from the scale forms (24). Indeed, all stem-chondrichthyan, non-acanthodian taxa in our dataset lack body fossils, while “Acanthodii” seems to represent a diagnostic set of ecologically-distinct lineages, or a grade of advanced “stem-Chondrichthyans” with both body fossils and diagnostic spines from throughout the Silurian-Permian (*2, 10, 16, 17*).

Given that the ongoing alternative use of “Acanthodii” and “stem-Chondrichthyes” in Ordovician-Silurian taxonomic surveys seems to be based on real differences in ecomorphology and ancestry, we have decided to assign our taxa to each following their attributions in the literature. That said, it is still possible that some of these distinctions are false given attribution based on different kinds of fossils. Further, it is possible that Ordovician scale-based “Chondrichthyes” taxa represent a distinct Ordovician lineage given differences with Silurian ichthyoliths assigned to the same group (see main text). To understand the influence of these classifications on our results, we have performed supplementary ecological reanalyses using three additional alternative groupings for fossils attributed to total group Chondrichthyes: 1) All Ordovician Chondrichthyes are “unknown”, Acanthodians and “Chondrichthyes” are separate Silurian groups, 2) All acanthodians and stem-chondrichthyans are “Chondrichthyes.”, and 3) Ordovician “Chondrichthyes” are unknown, all acanthodians are “Chondrichthyes.” The results of these alternative analyses are presented in the supplementary results (Figs. S10–16).

“Placodermi” is a category of stem-jawed gnathostome that has a similar history of usage to that for Acanthodii. Originally, Placodermi was considered to be a monophyletic clade outside the gnathostome crown, but is now thought to represent a polyphyletic grade of stem-lineages (e.g. Arthrodira, Antiarchi, Rhenanida, Ptyctodontida, Phyllolepida, and others) stretching from the origin of jaws to the origin of the crown (*2, 16, 48*). As placoderms have body plans that are conserved within their clades and distinct from most crown lineages, and have relatively few records in the Silurian, we have decided to treat their occurrences as belonging as a single group.

### Age and Environmental Assignments for Assemblages

Ages and geological stage assignments for our Ordovician-Silurian assemblages were determined based on the latest available data using the geological information within the original description or survey. Stratigraphic nomenclature and dates adhered to the “International Chronostratigraphic Chart” (2024/12) (*9*). All species within an assemblage were assigned the same age. Each assemblage was given a distinct name using its most commonly recognized designation, typically derived from a nearby settlement or landmark (Data. S1). In cases where multiple assemblages were confirmed in the same general area or from a continuous section representing different times, the name of the exact formation yielding the specimen. There were cases where the assemblage could not be assigned to a single stage, given boundary crossing faunas or uncertainty about exact location within a formation spanning multiple intervals.

Consequently, when calculating the total richness for each stage, the occurrences within these faunas were counted in multiple stages (Figs. 1, S3, S4; Tabs. S3, S5). However, when assigning faunas to a single time interval for the purposes of ordination, we chose the stage with the longer duration according to the “International Chronostratigraphic Chart” (*9*) (Figs. 2, 4A; Tabs. S1, S2).

The locality and geological information for assemblages was verified using the same approach as that for age determination, following Sallan and Coates 2010 (*16*). In addition, to interpret the distribution of Paleozoic gnathostomes, each fauna was assigned to one of 5 distinct geographic regions based on its locality: “Gondwana”, “Siberia”, “Laurasia”, “Baltic”, and “China” (Figs. 1–3, S4, S5C). These categorical region labels were subsequently converted to numeric values (1–5) for use in statistical analyses such as Canonical Correspondence Analysis and Non-Parametric Multidimensional Scaling (see below).

Evidence from geological literature or coincident invertebrate faunas was used to estimate the environment of our assemblages (Data S1). We used Benthic Assemblage Zones as a coding scheme spanning from freshwater (BA0) to open ocean (BA 6), following the procedure of Sallan et al. 2018 (*10*). The Benthic Assemblage zones for many of our assemblages were first determined by Boucot and Janis (*49*) or in Sallan et al., 2018 (*16*). For other assemblages without known habitat assignments, we referred to the literature as described in supplemental dataset (Data S1). After each fauna was assigned a Benthic Assemblage zone (Data S1), this was then simplified into three environmental categories: “freshwater”, “brackish” and “marine”.

Assemblages assigned as BA0 were treated as “freshwater”, while those where the fauna expanded into marine waters as well as BA0 were assigned as brackish. All other other BA values were treated as “marine”. These classifications were also subsequently converted into numeric values (1–3) for use in statistical analyses (Figs. 2B, 2C, S5B, S6, S11, S14; Tabs S7, S12). For some statistical analyses, the first two categories were combined in a single “non-marine” category by shape of plot or color (Figs. 2B, 2C, S5B).

### Datasets Used For Statistical Analyses of Faunal Composition

After compiling the raw occurrence and assemblage data in Data S3 and Data S4, the faunal details for each assemblage were compiled in Data S1, which represents our master list of Ordovician-Silurian faunas. Details include age, formation, locality, environment, the resident species assigned to each major gnathostome group, and sources (Data S1). Based on the age and taxonomic information for our assemblages, we first counted species-level occurrences within each stage, while special attention was given to assemblages that could not be precisely assigned to a single geological stage, either because of boundary crossing or lack of resolution for the exact age of the fossil-bearing units in the geological literature to date (Tab. S1). A total of 68 out of 168 assemblages were identified as covering multiple stages and are listed as such in Data S1. Species occurrences in multi-stage assemblages were counted separately in each relevant time interval for the purposes of analyses requiring time binning. Taxonomic richness at the genus level was also counted for each included time interval (Tab. S2); this data is highlighted in red in Data S3, and S4. The 68 multistage assemblages were counted as occurring within each of their stages to calculate the number of assemblages per stage in Tab. S3.

For analyses and comparisons of faunal composition, we first created a matrix with the assemblages as rows and the major gnathostome groups as columns. Due to the quality of the fossil record, we input the number of species within groups at each assemblage, rather than the absolute samples or occurrences within species as is more routine in ecological analyses (*16, 50*). In using this approach, we followed the justifications laid out by Sallan and Coates (*16*). For example, changes in the relative species richness within groups across assemblages is reflective of changes in the population levels and relative diversity of those groups given long enough time intervals (*16*). In addition, unlike for the invertebrate fossil record, workers do not routinely record abundances for vertebrate fossils at Paleozoic localities, and do not record each field sample as a separate collection (*2, 16*), making it difficult to reconstruct assemblages based on proportions or perform subsampling analyses to control for collection effort. Our faunal matrix of species within larger taxonomic groups at each assemblage is designated as Data S2.

Our faunal matrix served as the basis for generating histograms showing richness within major gnathostome groups in assemblages from the Ordovician to the Silurian periods. (Figs. 2A, 4A). Additionally, we created an expanded matrix containing variables such as age, environment, and regional information, which is presented as Data. S8, in order to perform ordination analyses and other statistical tests of the relationships between these variables and relative richness within gnathostome groups. This second matrix excludes assemblages from Data. S2 with less than three resident species to ensure robustness in the subsequent analysis, following the reasoning of Sallan and Coates (*16*). The matrix in Data S8 served as the basis for generating all of the multivariate ecological analyses conducted in this paper such as Shannon-Wiener index assessment, hierarchical clustering, canonical correspondence analysis (CCA), non-metric multidimensional scaling (NMDS), factor analysis (FA), one-tailed analysis of similarity (ANOSIM), and similarity percentage (SIMPER), described in detail below (Figs 2B, S5–16; Tabs S5–13) *16, 50*).

### Simple Visualization Methods

In this study, three basic data visualization methods were used to observe changes in species richness through time: stage-binned genus-level diversity curves (Figs. 1, 4B, 4C, S3– S4), species-level faunal histograms (Figs. 2A, 4A) and regional species-level pie charts (Fig. 3). All of these were generated based on the datasets provided in the supplementary materials (Tab. S1, S2; Data. S1–S7). Genus-level diversity curves were generated for the following groups across four comparisons: 1) Ordovician-Silurian gnathostomes (N=417) vs. conodonts (N=480) (Figs. 1A–B); 2) 13 groups of Ordovician-Silurian gnathostomes (Figs. 1C, S3); Ordovician-Silurian gnathostomes in 5 geographic regions (Figs. 1D, S4); 4) Silurian-Devonian jawless gnathostomes (5 groups) vs. jawed gnathostomes (4 groups) (Figs. 4B, 4C). We calculated taxonomic richness within each group in each stage at genus level to create the diversity curves (Tab. S3). For our conodont vs. gnathostome and jawed versus jawless comparisons, we also calculated per million-year diversity rate within bins by dividing the raw genus-level richness by the duration of each stratigraphic stage, as defined in the “International Commission on Stratigraphy” (Figs. 1B, 4C, S3C, S3D, S4C, S4D) (*9*). As above, in cases where the best age estimate for an assemblage was spanned multiple stages, resident species were counted once in each stage bin. In our diversity curve plots, we added black lines at the beginning and end of the Hirnantian to indicate mass extinction/glaciation pulses during Ordovician (*14*).

Two histograms were generated to estimate diversity trends across assemblages through time using our matrix of number of species within each of 13 groups at each assemblage (Fig. 2A), which we then recategorized into jawed and jawless gnathostomes (Fig. 4A). For the latter, species within the Unknown category were omitted. On the x-axis, all assemblages (N = 168) were arranged in loose chronological order based on their estimated ages (Data S1). Here, the assemblages spanning multiple stages were assigned to the one with the longer duration based on the “International Chronostratigraphic Chart” (9) while faunas of the same age were ordered alphabetically. The y-axis represented species-level richness within each assemblage (Range 1– 52) (Fig. 2A).

We used our assemblage data and matrix to create pie charts for species-level diversity within each stage for 5 regions in order to infer rough biogeographic patterns and dispersal (Fig. 3), and test hypotheses such as the “Out of Gondwana” model (*19, 51–53*). In our main figures, each chart was placed on a stage-specific paleomap (created by Ronald Blakey for “Deep Time Maps”) (*54*) near the center or main gnathostome fossil-bearing area of each paleocontinent, also representing the one of the five major Paleozoic regions described by Blakey (*54*).

### Multivariate Ecological Methods

To briefly compare the diversity of assemblages within each stage, three diversity indices (species count, Shannon-Wiener diversity, and taxonomic richness) were calculated based on species occurrences within stages and within faunas (Tab. S6) (*50, 55*). The Shannon-Wiener index assesses both richness and evenness within each fauna, while taxonomic richness does not consider evenness (*50*).

We visualized and analyzed the species and group composition of each assemblage using multivariate ecological methods and ordinations (*50*). As above, faunas containing only one or two species were omitted. The remaining dataset of 99 analyses was analyzed using R (version 4.4.2) (*56*) with the *vegan* package (version 2.6–10) (*57*) and PAST (version 1.0.6) (*58*). The applied multivariate methods included hierarchical clustering, canonical correspondence analysis (CCA), non-parametric multidimensional scaling (NMDS), factor analysis (FA), one-tailed analysis of similarity (ANOSIM), and similarity percentage (SIMPER) (*50*). All ecological analyses were performed using raw species counts or presence-absence within groups, as low absolute diversity for some assemblages made it difficult to use relative species diversity without introducing bias (Data S8).

To determine whether assemblages clustered by age, environment, or region, hierarchical clustering analysis was used to construct a dendrogram based on a distance or similarity matrix, without predefined grouping (*50*). The dendrograms were generated using the *hclust* function of stats package in R, employing the average linkage clustering (UPGMA) method (*59*). Clustering was based on Bray-Curtis distances calculated via the vegdist function in the vegan package (*50*, 57). Sites were color-coded according to age, environment, and region (Fig. S5).

Canonical Correspondence Analysis (CCA) allows us to visualize the distribution of faunas without requiring a priori grouping of sites (assemblages). This ordination analysis detects gradients in faunas based on associations between site composition and habitat or other variables such as time. It also allows taxonomic units and assemblages to be plotted together, showing the influence of specific groups on the distribution of sites (*50, 58*) Previous studies (*2, 10, 16*) have found a relationship between vertebrate assemblage composition and habitat depth, and shown that CCA analysis can detect significant differences between faunas across time.

Thus, two explanatory variables, geological series and aquatic environment, were used for CCA, which was generated by using *cca* function in the *vegan* package in R (Figs. 2B, S6, S10, S11, S14). In the taxon ordination plot (Fig. 2B, right), the direction of the two explanatory variables are shown as black dashed arrows. In our main figures, only the eight taxonomic groups with the highest contributions to the ordination were highlighted to enhance figure clarity, while all other taxa were omitted from the plot for simplicity.

A non-constrained ordination method, Non-Parametric Multidimensional Scaling (NMDS), was also used to visualize the distribution of assemblages, as this analysis does not assume a gradient or require environmental or other age variables unlike CCA. It also does not assume a normal distribution of samples unlike Principal Coordinate Analysis (PCA) (*50, 60*). This method visualizes the dataset in a reduced dimensional space while preserving the rank order relationships within the data, and is calculated from a dissimilarity matrix. Initially, Bray-Curtis abundance distance and Kulczynski presence-absence similarity metrics were applied to assess differences in the assemblage dataset spanning the Ordovician to Silurian periods (Figs. 2C, S7, S8, S10, S12, S13 S15, S16). In addition, comparisons were conducted between successive series intervals: Middle Ordovician and Upper Ordovician, Upper Ordovician and Llandovery (Silurian), Llandovery and Wenlock (Silurian), Wenlock and Ludlow (Silurian), and Ludlow and Pridoli (Silurian) (Fig. S7). These analyses were performed using the *metaMDS* function of the *vegan* package in R (58). The ordination was calculated in two dimensions (k = 2), with trymax set to 100 to ensure convergence to a stable solution. A random seed was set (set.seed(123)) to ensure reproducibility. Stress values were monitored to assess the quality of the ordination.

Factor Analysis (FA) was employed to identify key taxa contributing to faunal differentiation through visualization, and detect breaks across the mass extinction following prior usage by Raup and Sepkoski (*16, 50, 61*) (Figs S9, S10; Tab S8). This method allowed us to identify hidden common factors that drive the variation among observed taxa. By focusing on the covariance structure, FA highlights the underlying patterns that influence the data, rather than merely accounting for all the variance observed in the dataset (*50, 61*). The analysis followed a similar procedure to Principal Component Analysis (PCA), but with a fundamental distinction that will be discussed below. While PCA seeks to identify principal components that account for the maximum variance within a dataset by considering the total variance, FA is specifically designed to extract common factors based solely on the covariance structure among variables (*16, 50*). This approach enables us to unveil latent factors that may influence the distribution of taxa, helping to interpret the ecological processes driving community differentiation. They were generated by using PAST (*58*).

To determine if differences or similarities between assemblages from different geological series shown in our ordinations were significant, we used one-tailed Analysis of Similarity (ANOSIM) using pairwise comparisons of all series (*16, 50*). This analysis uses permutations of samples within two bins to determine an R statistic showing the degree of similarity or difference among the compared populations within those bins, and the significance of the same. We used 1-million permutations for each pairwise comparison between all gnathostome assemblages from each Ordovician-Silurian geological series, generating p-values (α = 0.05) and R statistics (Tab. S9). Similarity Percentage (SIMPER), which compares the relative contribution of taxonomic groups to differences between groups of sites, was also used to interpret the results of ANOSIM and to quantify the relative contribution of taxonomic groups to observed differences between intervals (Tabs. S10, S11). ANOSIM was generated by using the *anosim* function in the *vegan* package in R and SIMPER was generated by using PAST (*56–58*).

## Supporting information

Supplemental_manuscript_PDF

## Acknowledgments

We thank Erica Messics, Riley DeVito-Hurley, Ian Beyer, Daniel Sun, and Tyler Carlton for assistance with compiling occurrences. We also thank Ivan Sansom, Michael Coates, Martin Brazeau, and members of the Macroevolution Unit for preliminary discussions. We are grateful for the help and support about the figure design provided by Pavel Puchenkov in the section of Core Facilities at Okinawa Institute of Science and Technology Graduate University. Vertebrate reconstructions are by Nobu Tamura and used under a CC BY-SA license. Paleomaps are ©Deep Time Maps and used under license from Ron Blakey for this academic publication.

## Author contributions

Conceptualization: LS

Methodology: WH, LS

Investigation: WH

Visualization: WH, LS

Writing – original draft: WH, LS

Writing – review & editing: WH, LS

## Data and materials availability

All data, code, and results are available in the supplementary materials or on Dryad (temporary review link): http://datadryad.org/share/AOhNvKgGWKZHK2LZJ1Q8xi8pl2Z8V_gi4Dk-P4FjTd0.

## Supplementary Materials

Supplementary Text

Figs. S1 to S16

Tables S1 to S15

References (*62*–*117*)

Data S1 to S8

