## Supplemental_manuscript_PDF for "Mass Extinction Triggered the Early Radiations of Jawed Vertebrates and Relatives (Gnathostomes)"

**The PDF file includes:**

Supplementary Text
Figs. S1 to S16
Tables S1 to S13
References *62-117*

**Other Supplementary Materials for this manuscript include the following:**

Data S1 to S8

### Supplementary Text

Extinction-related turnover between conodonts and gnathostomes. In the Ordovician, conodonts (stem-cyclostomes) dominated a range of marine environments, including offshore settings, whereas gnathostomes were largely restricted to nearshore habitats (*4, 10, 14*). Conodonts were abundant and have served as important index fossils for biostratigraphic correlation and been used to define stages from the Tremadocian to the Hirnantian in various regions, including Baltica, Laurasia, Siberia, China, and Argentina (*62*). Based on "The Paleobiology Database" (*34*), as of late 2024, a total of 363 conodont genera has been reported from the Ordovician period (Figs. 1A–B; Data S6), compared to only 31 genera of gnathostomes from our more extensive dataset (Tab S2). In addition, previous work has found that conodonts occupied deeper waters in the Paleozoic and particularly in the Ordovician than gnathostomes, which exhibited environmental restriction nearshore (*10*). Despite both groups being affected by the mass extinction events of the Ordovician (14, main text), recovery of previous levels of diversity was confirmed only for Gnathostomes (Figs. 1A–B). Although gnathostomes were also impacted, as evidenced by the complete disappearance of Arandaspida and Astraspida after the Hirnantian (Figs. 1C, S3A–B), they recovered in diversity and eventually surpassed conodonts around the Aeronian stage of the Silurian (Figs. 1A–B).

Ordovician gnathostome occurrences. We confirmed the earliest known appearance of gnathostomes (*Porophoraspis sp.*) in the "Pacoota Sandstone" assemblage of Australia, which was the only site containing gnathostomes during Tremadocian (*63*) (Figs. 1C, 2A) (Data. S1–S3). Subsequently, members of the subclass Arandaspida (*Sacabambaspis sp.*), characterized by robust bony shields, were discovered around Gondwana in South America and Oman (*39, 64, 65*). Members of the Astraspis subclass, with similar ecomorphologies to Arandaspida, have been recovered from the Darriwilian of South America (*66, 67*), Iowa (*68*), Australia (*63*) and possibly in Siberia (*69*). Most of the Gnathostome records actually came from Gondwana until the end of this interval (Figs. 1D, 3A, 3B) (Tabs. S3A, S4A). While Arandaspida remained endemic to the Gondwana region, and recovered only from the outer regions (*70*), Astraspida gradually dispersed to other places, such as in Laurasia and Siberia region in the later Ordovician (Figs. 3B, 3C). The most abundant fauna of the Ordovician, "Canon city" (Shannon index: 1.28) appeared in North America during this time, and featured astraspids among other scale forms (Tab. S5; Data. S1). During the entire Ordovician, just 10 genera belonging to "armored" macromeric gnathostome classes (astraspids, arandaspids, and "pteraspidiformes") have been confirmed so far, and all were restricted to shallow marine environments at their localities (Data. S1) (*10*).

The only other positively identified Ordovician gnathostome groups are Chondrichthyes and Thelodonti, with taxonomic identification based only on scales as discussed above. Potentially chondrichthyan polyodontode scales (eg., *Tantalepis gatehousei*) have their earliest records in the Darriwilian of Australia (*71*). In the later Ordovician, chondrichthyan-like scales were recovered alongside with macromeric species such as *Astraspis*, and Acanthodii-like enigmatic ichthyoliths (*Skiichthys halsteadi*), but only at North American sites such as "Canon City" during Sandbian to Katian (*72–74*). Thelodonti, which seemed to be the most dominant gnathostome group in Silurian (Figs. 1C, 2A, 3) (Tab. S1, S2), first appear in Canada during Sandibian to Katian (*75*), and are also recovered from the Siberia region by the end of the Ordovician. However, as mentioned above, the specimens of Thelodonti were still limited to rare and isolated cales and no macrofossil records have been discovered so far during the Ordovician.

Faunal and biogeographical changes across the LOME interval. There are significantly more marine than non-marine localities for Ordovician-Silurian gnathostomes. Out of 168 total assemblages, 30 were categorized as having "non-marine" environments, including 6 in "freshwater" and 24 in "brackish water" (freshwater influenced, or euryhaline) settings (Data S1). Only two of these non-marine or brackish faunas, "Split Rock" and "Canon City", belonged to the Ordovician; the remaining 38 occurred during the Silurian (Data S1). Among the non-marine faunas, 18 had more than two taxonomic occurrences and so could be used for our ordinations and other analyses, compared to 81 such faunas in marine settings. There is not much difference between the composition of marine and non-marine assemblages from the same region in our ordinations (Figs. 2A, 2C). We recovered 37 Ordovician gnathostome assemblages and 203 Silurian assemblages (Tab. S4). Until the Darriwilian, most of these assemblages (14 out of 16) had occurred in the Gondwana region such as Australia, South America and Arabian peninsula, however a few faunas ("Angara River" and "Winneshiek Lagerstatte") were also recovered in Siberia and Laurasia regions outside of Gondwana (Tab. S4; Data S1). In the Sandbian, the number of gnathostome localities in Laurasia overtook that of Gondwana, and most Gnathostomes including possible chondrichthyans and other jawed fishes were discovered in this region (Fig. 1E; Tab S4). During the Hirnantian stage, when a glaciation event occurred at the South Pole in Gondwana (*76*), both Gondwanan and Laurasian gnathostome faunas disappeared, with only a few low-diversity assemblages (e.g., "Thunia River") recorded in the Siberian region remote from Gondwana.

Following the Ordovician, Siberian faunas continued to be documented consistently until the end of the Silurian, and a Laurasia fauna also appearing continuously after the Rhuddanian. In contrast, the appearance of new Gondwanan faunas was delayed, first emerging in the Homerian, approximately 15 million years after the extinction. This is with the exception of the South China area, which was isolated from the South Pole in Gondwana by both the Paleotethys Ocean and the Panthalassic Ocean (*77, 78*). In the Telychian, the total number of sampled gnathostome assemblages reached its Silurian peak (N = 33), even though global genus richness did not peak until the late Silurian, suggesting lower average per-assemblage richness and homogeneity within regions (Figs. 1A, 1B, 2A). During the Telychian, the Chinese fauna (9 out of 33 assemblages) was composed of unique species such as jawed gnathostomes and galeaspids (Fig. S5C), as explained below and in the main text (Fig. 3D). Many of these assemblages were previously regarded as Wenlock in prior work (*79*), however they have been re-assigned as Llandovery by recent research (*80*).

Gnathostome victims and survivors of the LOME. Multiple ecological and environmental changes are associated with the polar glaciation pulses in Gondwana at the end of the Katian and Hirnantian (*14*). These events were marked by falling sea levels, which caused the marine ecosystem to totally collapse, particularly in offshore areas (*81*). In our figures, we show mark these glaciation pulses to show the timing of extinction events in relation to vertebrate diversity trends (*14*) (Figs. 1A–B, 2A, 4A).

Some Ordovician gnathostomes were affected by these events as discussed in the main text, while others seemed to survive and diversify during the recovery interval (Figs. 1C, 2A, S3). After the first extinction pulse, uniquely Ordovician taxa such as Astraspida and Arandaspida completely disappeared, which is the source of their significant contribution to the negative region of the geological time (series) correlated axis 1 in our CCA (Fig 1C, 2B; Tab.

S5). On the other hand, some Ordovician thelodonti, such as Sandiviiformes (e.g., *Larolepis darbyi*) which appeared in Laurasia during Sandbian (*72*) (Fig. 3C; Data. S2, S3), seemed to have survived the initial pulse of the LOME in Siberian refugia where they dispersed during Katian (*81*). Although these specific thelodonts lineages were replaced in the earliest Silurian by Loganelliiformes in Siberia (*Loganellia, Talimaalepis*), Thelodonti dispersed widely after the end-Hirnantian and showed the highest contribution to our Factor Analysis (Factor1: 3.60) during Silurian (Figs. 2A, 3; Tab S8) (*82, 83*).

Among other Ordovician gnathostomes, potential chondrichthyan or jawed gnathostome scale-based taxa (eg., *Tantalepis gatehousei*) disappeared from Laurasia region before the beginning of the extinction interval, and a new elegestolepid group appeared in the Siberian fauna ("Moyerochan Formation") beginning in the earliest Silurian (*83*) (Figs. 1A, 3D; Data. S1). Elegestolepid and Mongolepid ichthyoliths continued to survive in the same region (eg., "Elegest and Kadvoi2") until Sheinwoodian (Figs. 1A, 3E; Data. S1). In addition, a new type of related chondrichthyan ichthyolith, *Qianodus duplicis,* bearing the possible earliest evidence teeth, suddenly appeared at 439 Ma in the Chinese region (*84*). The first body fossils of total group Chondrichthyes (the acanthodian grade *Shenacanthus vermiformis*) have been confirmed from the same time and region (*8, 24*) (Fig. 2A; Data. S1). However, no tooth or jaw material similar to these confirmed chondrichthyans has been recovered from the Ordovician or earlier Silurian, and there remains a gap in the potential jawed gnathostome record between the Katian and early Llandovery (Figs. 1C, 2A; Data S1).

<u>The first appearances of gnathostome lineages in the Silurian.</u> Many new gnathostome lineages and classes appeared after the LOME, at the same time that global gnathostome richness was drastically increasing in the Silurian (Fig. 1C). Except for Osteichthyes, all Silurian gnathostome classes described appeared by the Telychian, less than 10 million years after the second extinction pulse (Fig. 1A). Distinct clades of macromeric jawless gnathostomes such as Osteostraci (e. g., *Kalanaspis delectabilis*), Galeaspidomorphi (*Jiangxialepis retrospina*)*,* and Heterostraci (e. g., *Ariaspis sp.*) appeared in the Llandovery Telychian of the Baltic, China and Laurasian region, respectively (Fig. 3D) (*85–87*). Osteostraci and Heterostraci dispersed throughout by the end of the Silurian to all continents with the exception of Gondwana and China, whereas Galeaspidomorphi remained endemic to the China region throughout the Silurian (Fig. 3E–3G). Among these groups, Heterostraci showed the highest contribution to the distribution of assemblages in our Factor Analysis (Factor 3: 3.59) and their taxonomic richness of became highest by the Pridoli (Tabs. S2B, S2D, S8). A new micromeric jawless gnathostome group, Anaspida, had dispersed not only across Baltica but also into Laurasia and the Siberian regions alongside Thelodonti by the Wenlock, much earlier than other jawless groups. Their first appearance was in the Telychian-aged "Shank's Castle" locality (Fig. 3E) (*88*).

Among the jawed groups besides "Chondrichthyes", Acanthodii (*Tchunacanthus obruchevi*) appeared first in Russian faunas ("Kyzyl-Tchiraa" and "Moyerochan Formation") from the Rhuddanian (Figs. 1A, S3B, S3D; Data. S1). They appear to have successfully dispersed globally by the end of the Silurian (Fig. 3). Placodermi, which would become the dominant set of lineages in Devonian (*16*) is confirmed from Telychian based on well-preserved fossils (*Xiushanosteus mirabilis*) from China, which occurred alongside the whole body Chondrichthyes mentioned above (Fig. 3D) (*8*). However, placoderms remained endemic to the China region and restricted to the shallow water areas during the rest of the Silurian (Data. S1) (*10*). One lineage of Osteichthyes (*Andreolepis hedei*) finally appeared in the Baltic region around the Ludfordian (e.g., "Gotland3"), before their emergence in much larger numbers in China following a gap in the record of that region (Fig. 3F) (*89*). More specific lineages of osteichthyans, such as Sarcopterygii (eg., *Guiyu oneiros*), appeared in China around the same time, starting with "East Yunnan" suggesting diversification in that region (*90*).

<u>Changes in faunal composition over the LOME and Silurian based on multivariate analyses.</u> The composition of gnathostome biodiversity and assemblages species showed clear differences in between the Ordovician and Silurian in simple visualizations of our raw data as above. These differences were confirmed through our ordinations and permutation tests, as well as the influence of specific groups.

Canonical correspondence analysis (CCA), conducted with 101 fauna input and two explanatory values (Series and Environment), clearly separated all Ordovician and Silurian assemblages along axis 1, and all Ordovician and Llandovery assemblages on axis 2 (Fig. 2B, left). In the ordination plot showing the influence of variables and taxa on assemblage position (Fig. 2B, right), the arrow representing Series, or geological time, aligned closely with axis 1, which means the gradient of the time interval in each fauna was shown from left to right. Axis 1 accounted for over 90% of the total variance, highlighting time, or position before or after the LOME, as the dominant factor in differentiating between faunas. Our main figure 2B showed the 8 most important groups; all others generally fell between the arrow of Osteichthyes and axis 1 in the positive side (Fig. S6). Consistent with the simple visualization analysis shown above and the position of Ordovician assemblages, the arrows on the CCA plot positioned Ordovician-unique taxa on the negative end of axis 1. Among them, Arandaspida showed the lowest score along axis 1 (Tab. S7), supporting its strong association with Ordovician faunas and complete disappearance after these extinction events.

In terms of the CCA distribution of assemblages within each time bin, "Canon City" shown as "*F1*" in Upper Ordovician was the closest to Llandovery Silurian faunas on axis 1 (Fig. 2B, left and 2C). This proximity is likely due to the fact that it represents the only Ordovician fauna yielding fossil records for presumed Chondrichthyes. This is the source of ecological affinity with Chinese faunas from the Llandovery and later Silurian, which were tightly clustered on axis 1 between the Ordovician samples and Silurian assemblages from the rest of the world. However, there was a wider separation between Ordovician and Chinese faunas on axis 2, with those from elsewhere in the Ordovician falling between (Fig. 2B). Most of the Chinese faunas were located on the positive side of axis 2, with the brackish site "Bachu-Kalpin2" being the most positive. These Chinese sites are considered unique, as many represent freshwater or brackish environments (Fig. S5C) and yielded distinctive taxa such as Galeaspidomorphi and Placodermi during the Silurian which do not appear in the Ordovician or outside China in the Silurian. The biplot arrow representing the variable Environment was oriented somewhat in line with axis 2 but with limited effect, where the positive end of axis 2 reflects shallow-water settings based on the positive values of Chinese non-marine habitats indicated by the filled triangle symbols in the plot (Fig. 2B). However, this alignment may be a distortion caused by differences between Ordovician and Silurian faunas, as most non-marine and marine Silurian assemblages from the same region were overlapped while the only freshwater influenced Ordovician site ("Canon City") also had chondrichthyan remains (Fig. 2A).

In the NMDS analyses based on both Bray–Curtis (raw species counts) and Kulczynski (presence-absence of group) indices, the stress values across intervals ranged from 0 to 0.12, which was near or within the generally accepted threshold of 0.1 for reliable ordination (Figs. 2C and S7–8) (*60*). The NMDS plots separated the assemblages into Ordovician and Silurian groups, both across the entire dataset and especially when limited to the Upper Ordovician–Llandovery interval, although this separation was not aligned with any specific axis as in the CCA analysis (Figs. 2B, 2C). Compared to the Bray Curtis-based NMDS plot (Fig. 2C), the Kulczynski-based plot (Fig. S8) revealed a clearer separation. As in the CCA, "Canon City" (*F1*) was again the closest Ordovician fauna to the Silurian cluster. Several faunas, *F1–F3*, *F5*, *F6*, and *F8*, (Fig. 2C) showing maximum or minimum NMDS scores along the axes, overlapped with those identified in CCA (*F1–F4, F7*, *F8*) (Fig. 2B), indicating consistency between the two methods despite their different approaches.

As described in our methods, we used Analysis of Similarity, a permutation test, to determine the exact degree and significance of observed similarity and dissimilarity between sampled diversity from assemblages in different time intervals (*16*). This also confirmed a clear difference in species composition between Upper Ordovician and Llandovery (ANOSIM: mean $R$ =0.38, $p$ = 0.02) (Tab. S9). This result was further supported by Similarity Percentage (SIMPER) analysis, which revealed a 90.58% dissimilarity between the two intervals (Tab. S10). Nearly half of this dissimilarity was explained by differences in the presence of *Thelodonti* and *Astraspida*, taxa that were abundant in the Silurian and confined to the Ordovician, respectively.

The difference in species composition between the Ordovician and Silurian was also supported by Cluster Analysis (Fig. S5A). All Ordovician faunas were clustered on the lower part of the dendrogram, with Llandovery faunas separate. As shown in the CCA and NMDS results, the Silurian faunas, especially those from later stages, were more intermixed, suggesting homogeneity during that period. Also, in line with the overall CCA results, no clear clustering pattern was observed based on environmental values (Fig. S5B). In the CCA biplot (Fig. 2B, right), the arrow representing the "environment" variable was shorter than that of "series" and was aligned with axis 2, which accounted for substantially less variance than axis 1. Although environmental factors, particularly habitat depth, have been previously shown to influence species composition (*10, 16*), our results suggested that it was less of a factor in the distribution of major groups or the overall composition of assemblages, perhaps because so few lineages were exclusively freshwater or deep water throughout most of this period. When we color coded the assemblages in our cluster analysis by region (Fig. S5C), clear clustering patterns were observed, especially among Chinese faunas (e.g., "Qingshui Formation") characterized by endemic taxa. In fact, the cluster of early-mid Silurian Chinese faunas represented the outgroup to all other assemblages. However, later Silurian Chinese faunas, such as "East Yunnan", were mixed in with those from other regions. This reflects the invasion of thelodonts into Chinese faunas in Ludlow and greater dispersal of some jawed fishes during the same period (Fig. 3E). In addition, Laurasian faunas formed a distinct cluster near the bottom of the dendrogram. This cluster yielded fossil records of Heterostraci and included six faunas—"Somerset Island3," "Yukon," "Leopold Formation," "Yukon2," and "Avalanche Lake"—which represented the sites with the highest number of Heterostraci specimens discovered in the Silurian (Data S4).

Factor analysis appeared to be less effective in capturing the compositional differences between Upper Ordovician and Silurian faunas (Fig. S9). The factor scores for Ordovician-unique taxa such as *Astraspida* were relatively low (Factor 1: 0.04, Factor 2: –0.16), compared to those of other groups that contributed more strongly to the ordination. In particular, *Thelodonti* showed a high score along Factor 1 (3.59), while *Chondrichthyes* and *Galeaspida* showed strong negative scores along Factor 2 (–2.34 and –2.72, respectively), suggesting that the variation captured by the analysis primarily reflects Silurian taxa. This may be partly due to the simplified classification scheme used in this study, in which only three environmental categories were defined, reflecting the difficulty of accurately assigning single benthic assemblage (BA) zones for each assemblage.

Comparison of Silurian-Devonian diversification in jawless gnathostomes and jawed gnathostomes. We merged our Devonian genus-level occurrence dataset (Data. S5) with our occurrences for the Silurian in order to determine longer term trends following the recovery interval. Jawless gnathostomes showed much greater global richness from Ordovician to early Devonian, even after definitive jawed gnathostome body fossils appeared in the Silurian, and after suspected chondrichthyans in the Ordovician (Fig. 4). The most popular hypothesis explaining the extinction of jawless fishes is that jaws were a key innovation that allowed filling of new niche space and competitive exclusion of jawless fishes (*2, 10, 45*). An alternative hypothesis suggests that jawless fish diversity was related to sea level, and therefore these lineages more restricted by habitat and mobility than jawed fishes (*12, 91*). However, according to our dataset, it took over 60 million years for jawed fishes to surpass jawless gnathostomes in global richness (Figs. 4B, 4C), perhaps because of the restriction of the latter to China for most of the Silurian and slow dispersal thereafter (Fig. 3) (*10*). Both of these groups had experienced multiple diversification pulses within restricted regions during Ordovician to Devonian (*10*), but the appearance of major jawed gnathostome lineages, and their dispersal, was slightly delayed to those of even shallow water benthic jawless gnathostomes (Figs. 3, 4B, 4C). Jawed and jawless gnathostomes exhibited coincident diversification pulses from the Pridoli to the Pragian in terms of genera per million years (Fig. 4C). This followed the end of the Silurian recovery interval (*15*) and their dispersal of shallow water forms out of their initial ranges (*10*). Jawless gnathostome global then drastically declined over the Emsian-Eifelian, in line with an increase in jawless fish richness per stage (Fig. 4B), or at least a less significant faltering when time corrected (Fig. 4C). These lineages never recovered; jawless fishes were only a minor part of vertebrate faunas after the mid-Devonian, and every major group went extinct before the end of the period (*2, 10, 17*)(Fig. 4B, 4C). In contrast, jawless fishes reached peak stage-level global richness in the late Devonian (Fig. 4B), at the same time they achieved maximum dispersal in terms of habitat and most major lineages became cosmopolitan (*2, 10, 16, 17*). However, our time corrected diversity curve suggests that jawed lineages never achieved the global peaks in genus richness exhibited by the jawless fishes in the early Devonian (Fig. 4C).

The global extent of 'Talimaa's Gap' and regional intervals of low post-extinction diversity. Our dataset confirmed a previously proposed 3–5million year interval of low and absent diversity in the gnathostome global and regional records between the Ordovician and early Silurian, a period known as "Talimaa's Gap" (*2, 21, 22, 10*). Following the Hirnantian glaciation event around Gondwana, we did not find published evidence of any gnathostome assemblages in most of Gondwana until late Homerian ("Derenjal Mountains") (Figs. 1D, 3E) (Data. S3, S4) (*92*). The exception was China, then part of the equatorial archipelago at the margin of Gondwana, where new gnathostome faunas suddenly appeared at the beginning of the Silurian (Figs. 1C, 2A, 3D). This region yielded faunas with unique taxa such as Galeaspidomorphi, which emerged abruptly during the Llandovery (Figs. 1C, 2A, 3D), despite the continued absence of any Chinese records in the Ordovician or bridging the Ordovician–Llandovery boundary. Although 9 out of the total 18 known Chinese Silurian assemblages date to the Telychian Llandovery, the number of records declined afterward and did not increase again until the Ludfordian (Fig. S3). One of the reasons is that recent stratigraphic revisions have reassigned many mid-Silurian faunas to older ages than previously estimated shifting several sites once considered Wenlockian into the Llandovery interval (Data. S1) (*90, 93, 94*).

Looking to other regions, a low number of assemblages were consistently recorded in Siberia from the Hirnantian to the Rhuddanian (Data S1) (*39, 82, 83, 95*). In contrast, only one occurrence ("Petit River Port Daniel") was documented in Laurasia during the same interval (*96*). While Siberia yielded rich assemblages with a diversity of gnathostome groups from the earliest Silurian (e. g., "Moyerochan Formation"), Laurasian sites with at least 3 resident gnathostome taxa did not appear until the Telychian ("Devon Island") (Fig. 2A). These regional differences likely reflect the uneven influence of environmental changes during the Hirnantian glaciation, as these were interpreted as discontinuous across Gondwana (*97*). The delayed recovery in Laurasia may be attributed to harsher post-extinction environmental conditions or to the relative absence of suitable refugia during the early Silurian.

Ordovician-Silurian gnathostome refugia and later Silurian biogeography. There was some evidence for specific refugia for boundary-crossing gnathostomes in different regions during the Ordovician-Silurian. In equatorial Siberia, a few gnathostomes seemed to have persisted during and after the Ordovician mass extinction events at high enough richness to be captured by the record (Fig. 2D), perhaps due to being remote from the ice cap region (*97*). The Shannon diversity index of the "Moyero River" fauna, one of the earliest Silurian assemblages in the Siberian region, was comparably high to that of "Canon City," the most diverse Ordovician fauna (Moyero River: 1.28, Canon City: 1.28). This high diversity suggests that Siberia might have functioned as a refugium during the extinction crisis (Tab. S6) (*49, 74, 83*).

Similarly, the China region, where new faunas emerged at the onset of the Silurian, might also have acted as a refugium for early gnathostomes. Although this region was surrounded by both the Paleo-Tethys Ocean and the Panthalassic Ocean, it consisted mainly of shallow-water environments along the margin of Gondwana (*98, 99*). These geographic features might have facilitated the inflow of species from the Gondwanan shallow seas while restricting their dispersal to other regions, thereby allowing unique lineages to persist and evolve in isolation. In fact, Ordovician brachiopods of the Kazakh terranes were confirmed to disperse between South China and Australia (*99*).

The Baltic region also played a key role as a refugium in the Silurian. It was here that the earliest Osteostraci (*Kalanaspis delectabilis*) appeared (*85*). This region shared several Thelodonti taxa with Siberia and Laurasia, suggesting possible migration from Siberian refugia (Data S1). In Laurasia, Ordovician macromeric taxa such as Astraspida were replaced in the Silurian by Heterostraci, which share some key traits (see methods) and appear to have occupied similar ecological niches (*10*) (Fig. 3). Heterostracans subsequently dispersed into Siberia and the Baltic regions (Fig. 3). In parallel, Osteostraci became increasingly diverse, especially in the Baltic region from the Wenlock onward. It is important to note 19 out of the total 30 Silurian non-marine assemblages were discovered in the Laurasian and Baltic regions, and that these also contain specific jawless lineages. This pattern may reflect the development of mountain belts in these regions, which brought continental landmasses closer together through plate tectonic processes, thereby facilitating the expansion of freshwater and brackish habitats suitable for non-marine vertebrates (*100*).

Prolonged regional isolation of jawed gnathostomes. As noted above, well-preserved jawed gnathostome records were recovered from China starting in the early Silurian (Fig. 3D). In "Chongqing," the earliest body fossils of "acanthodian-grade" chondrichthyans and placoderms, the latter of which dominate Devonian oceans (*16*), were found alongside Galeaspidomorphi (*8, 28*) (Fig. 2A; Data. S1). Osteichthyes did not appear in South China until the later Silurian, dating after isolated elements from the Baltic region and a hiatus in the Chinese record (Fig. 1C), but demonstrated high richness within Chinese assemblages (e.g., "East Yunnan") (*79, 101, 102*). This suggests that Silurian Osteichthyans also originated in China, in line with evidence from *Entelognathus* and later stem members of the clade (2), while those outside are early dispersers. In fact, by the late Silurian, plate movements brought the Chinese blocks closer to Siberia, Laurasia, and the Baltic (*103*) in line with nascent jawed fish dispersal across the broader region and allowing invasion by outside lineages such as thelodonts (Fig. 2G). This tectonic shift might precede the increase in global jawed gnathostome richness observed in our diversity curves (Figs. 4B, 4C)

Biases in the thelodont microfossil record. Thelodonti was the most dominant taxa in our global and assemblage-level datasets and showed a significant impact on several statistical analyses, as one of the few lineages to cross the extinction boundary unchanged in distribution or basic form (Figs. 3C, 3D; Tabs. S1, S2, S5, S6, S9). However, most of their Silurian record and taxonomy was based on microfossils such as isolated scales; no macrofossil evidence existed for Thelodonti before body fossils for *Loganellia* in the Telychian fauna of "Lesmahagow" (*104, 105*). While this has historically raised concerns about taxonomic resolution, scale-based classification in thelodonts is widely accepted and supported by phylogenetic analyses (*106, 107*). Previous research has also suggested that surveys and published records of thelodonts morphological diversity is still incomplete (*108*), implying that many taxa might remained undescribed and that we have underestimated known thelodont diversity. In addition, although the affinity has been proposed between Sandiviiformes (Ordovician Thelodonti group) and Loganelliiformes (Silurian dominant Thelodonti group) on the basis of scale histology and form (*109*), major differences have also been noted (*71*). Moreover potential affinity between various Sandiviiformes and other taxa such as Chondrichthyes and Pteraspidomorpha was also porposed (*23, 110*) suggesting this groups might not be monophyletic. These perspectives raise the possibility that early "Thelodonts" may eventually be split into multiple class-level lineages, supplying a richer and more complex picture of early vertebrate evolution in Ordovician and Silurian.

Uncertainty in the microfossil record of Chondrichthyes. Like thelodonts, the fossil record for Ordovician-Silurian Chondrichthyes was also dominated by microfossil-based taxa, as discussed in the methods section. The earliest known stem-Chondrichthyan body fossil, *Fanjingshania*, appears in the late Aeronian in Chinese fauna, "Leijiatun Village" (*24*)(Data S1). This preservational gap following the first appearance of possible Chondrichthyan scales in the Ordovician is consistent with other recent independent compilations of the record (*111*). We also discovered another gap in the overall stem-chondrichthyan record during the middle to late Silurian; after the appearance of *Elegestolepis grossi* and *Udalepis* sp. in the Wenlockian Siberian fauna, "Elegest and Kadovoi 2" (Fig. S3) (*112*) no records have been recovered older than an indeterminate specimen from the Laurasian region, "Hall land2" (*113*).

Our data suggest that gaps in the Chondrichthyan microfossil record during the mid-Paleozoic may be attributed to two causes. First, Ordovician-Silurian chondrichthyan form taxonomy is still controversial, with much uncertainty among even expert workers (*2, 3, 10*). As noted in our methods, we grouped most Ordovician taxa attributed to Chondrichthyes into the "Unknown" category based on alternative published assignments and uncertainty expressed within the descriptions, with the exception of well-described specimens from "Canon City" which have been consistently assigned as Chondrichthyes by the same authors (*38*). Aside from the well-preserved early Silurian teeth and body fossils from China, *Elegestolepis* microfossils from the Moyero River in Russia are the oldest known Silurian record for non-acanthodian Chondrichthyes, however these also showed affinity with Thelodonti (*23, 83*). Spines from the same assemblage, *Tchunacanthus* and *Lenacanthus* have also been attributed to "Acanthodes", although their exact taxonomy still remains unresolved (*114).*

Several studies have proposed that Acanthodian taxa such as *Climatiiformes* and *Ischnacanthus*, which were confirmed in the early Silurian and were categorized as "Acanthodii" in this study, represent a paraphyletic stem groups of Chondrichthyes (*2, 3, 115–117*). If these Acanthodian lineages are included Chondrichthyes, they may partially fill the aforementioned gaps. Our data suggest more complete and diagnostically Devonian-like chondrichthyan material tends to be assigned as "Acanthodian" rather than "Chondrichthyes", including the earliest body fossil which was described as "acanthodian grade" (*24*). Indeed, by the late Silurian, non-acanthodian stem-Chondrichthyans disappeared almost entirely from our assemblages (Fig. 2A), and our CCA results suggest that the taxonomic use of "Chondrichthyes" rather than "Acanthodii" was particularly associated with scale and spine forms at early Silurian sites (Fig. 2B). Therefore, while there was a shorter gap between the Wenlock "Chondrichthyes" and Ludlow acanthodians, the Chondrichthyes-specific mid-Silurian gap might be a result of an increase in diagnostic stem-chondrichthyan material and body fossils at later Silurian assemblages.

Effects of reclassifying Ordovician Chondrichthyes as "Unknown" on faunal multivariate analyses (Reanalysis 1). To determine the influence of chondrichthyan taxonomic practices on our results, we repeated the ordination and statistical analyses of our assemblages using alternative groupings for Chondrichthyes and Acanthodii (see methods for explanation). As noted, "Canon City" is the only Ordovician assemblage that has yielded fossil scales attributed to Chondrichthyes without any published taxonomic alternatives or disputes involving the original authors. In our main ordinations, "Canon City" was positioned closer to Llandovery assemblages than other Ordovician faunas, and even overlapped on some higher axes (Figs. 2B, 2C, S6–9). Given the fragmentary nature of supposed jawed gnathostome records in the Ordovician and the possibility of boundary crossers among low diversity ghost lineages (*2, 3, 10*), we wanted to determine how attribution of Ordovician scale taxa to crown jawed lineage influenced out results.

We performed analyses of an alternative dataset in which all Ordovician Chondrichthyes, including those from "Canon City", were reclassified under the "Unknown" category (Fig S10; Tab S12). In the reanalysis, the overall pattern of the CCA remained broadly consistent with the main results (Figs. 2B, S10), although the variation explained by axis 1 (time) was slightly higher (Tabs S7, S12). The faunas with the highest and lowest scores along each axis (*F1–F9*), as well as the top eight taxa contributing most to the ordination, remained largely unchanged between the analyses. In contrast, more substantial differences were observed in NMDS and FA when focusing specifically on the interval spanning the Upper Ordovician to the Llandovery (Figs. 2C, S10). In both analyses, the separation between these two stages became more pronounced in the reanalysis. Thus, the apparent close affinity of "Canon City" with Silurian assemblages is likely dependent on the classification of scales as belonging to crown gnathostomes, as the rest of the astrapid-dominated fauna is more in line with other Ordovician assemblages (Fig. 2A).

Effects of reclassifying all Acanthodii as Chondrichthyes on faunal multivariate analyses (Reanalysis 2). To evaluate the impact of treating Acanthodii and other stem-Chondrichthyes (not designated at Acanthodii in the literature) as separate groups, we conducted an alternative analysis in which all Acanthodii were grouped with Chondrichthyes (Figs. S11–13; Tab S12). In the CCA (Fig. S11), the left panel shows that faunal separation between the Ordovician and Silurian was retained, but the explanatory power of Axis 1 (time) was slightly reduced compared to the main analysis (Fig. 2B). In the right panel, the taxon arrow for Chondrichthyes was replaced by Heterostraci, despite the continued inclusion of Ordovician Chondrichthyes. This likely reflects the increased spatial and temporal distribution of Chondrichthyes resulting from the inclusion of globally dispersed Acanthodii (Fig. 3). In the NMDS ordinations (Figs. S12, S13), the separation between Ordovician and Silurian faunas became more distinct under both Bray–Curtis and Kulczynski distance metrics, relative to the main analysis (Figs. 2C, S7, S8). Notably, the position of the "Canon City" assemblage, which previously clustered near early Silurian faunas due to the presence of Chondrichthyes, shifted toward the Ordovician group in this reanalysis. This reflects the reduced relative contribution of Chondrichthyes in that assemblage once the group was redefined as widespread. Despite these changes, the position of other early Silurian faunas remained stable, supporting the idea that post-extinction faunal turnover was primarily driven by genuine ecological reorganization rather than taxonomic misclassification. This was expected given the occurrence of "Chondrichthyes" on both sides of the extinction boundary in the main analyses (Figs. 1A, 2A).

Effects of reclassifying Ordovician Chondrichthyes as "Unknown" and Silurian Acanthodii as Chondrichthyes on faunal multivariate analyses (Reanalysis 3). Finally, we conducted a reanalysis in which all Ordovician Chondrichthyes were reassigned to the "Unknown" category, while all Acanthodii were reclassified as Chondrichthyes, incorporating both of the above conditions (Figs. S14–S16; Table S12). The overall pattern in both CCA and NMDS plots remained consistent with the main analysis, but the separation between Ordovician and Silurian faunas became more pronounced (Figs. 2B–C). In the right panel of the CCA (Fig. S14), the arrow representing Chondrichthyes was again replaced by Heterostraci, and the vector for "Unknown" became longer than in the main result, indicating increased influence of unclassified early taxa.

These results reinforce the sensitivity of multivariate analyses to taxonomic treatment at the boundary interval. While adjustments to classification can influence the placement of specific assemblages—especially those with ambiguous early gnathostome material—the overarching structure of faunal turnover remains robust. As in the previous reanalysis, the stability of early Silurian assemblage positions supports the conclusion that the Ordovician–Silurian transition reflects true ecological reorganization rather than an artifact of inconsistent taxonomic assignment.

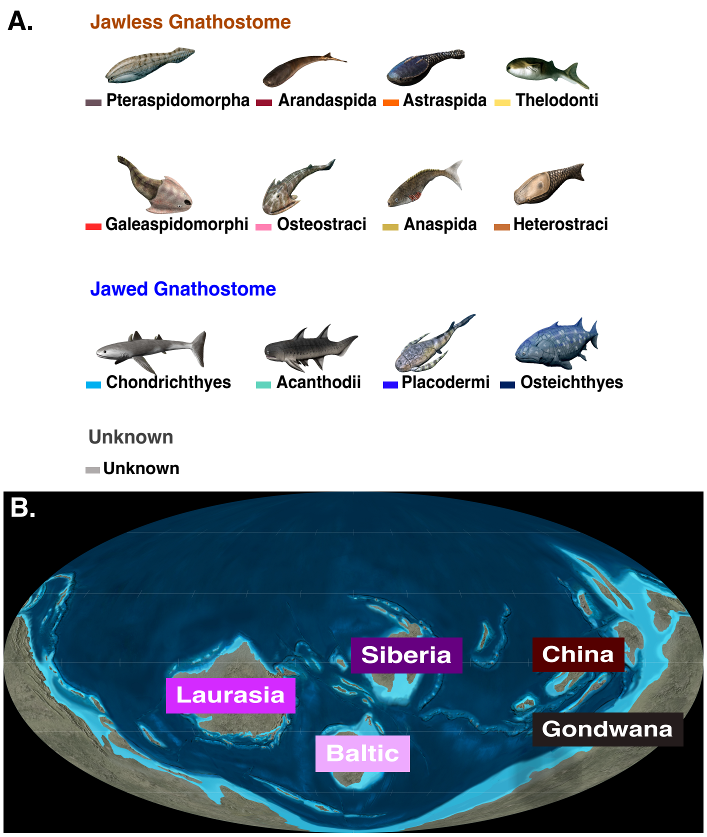


**Fig. S1. Paleozoic gnathostome classes and regional map from the Ordovician to Silurian.** **(A)** 13 classes of Paleozoic gnathostomes, each represented in a different color. Eight classes were assigned as **Jawless Gnathostomes**, and four classes were assigned as **Jawed Gnathostomes.** See Material and Methods for explanation. **(B)** The 5 distinct regions where gnathostome assemblages were recovered. Paleomap from Deep Time Maps (*54*) and used with permission. Colors for groups and regions were used in Figs. 1C, 1D, 2, 3, S4 and S5.

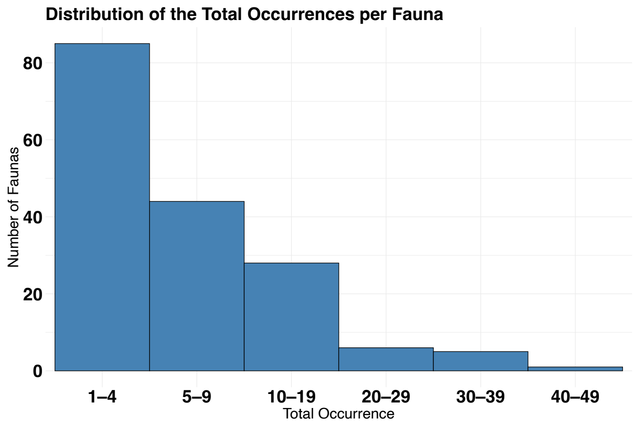


**Fig. S2. Histogram of species-level diversity at Ordovician-Silurian gnathostome faunas (assemblages).** Total Occurrences = 1,156; Total Faunas = 168; Mean of Occurrences = 6.85, SD of Occurrences = 8.11, Median of Occurrences = 4. See Data S1 for more details.

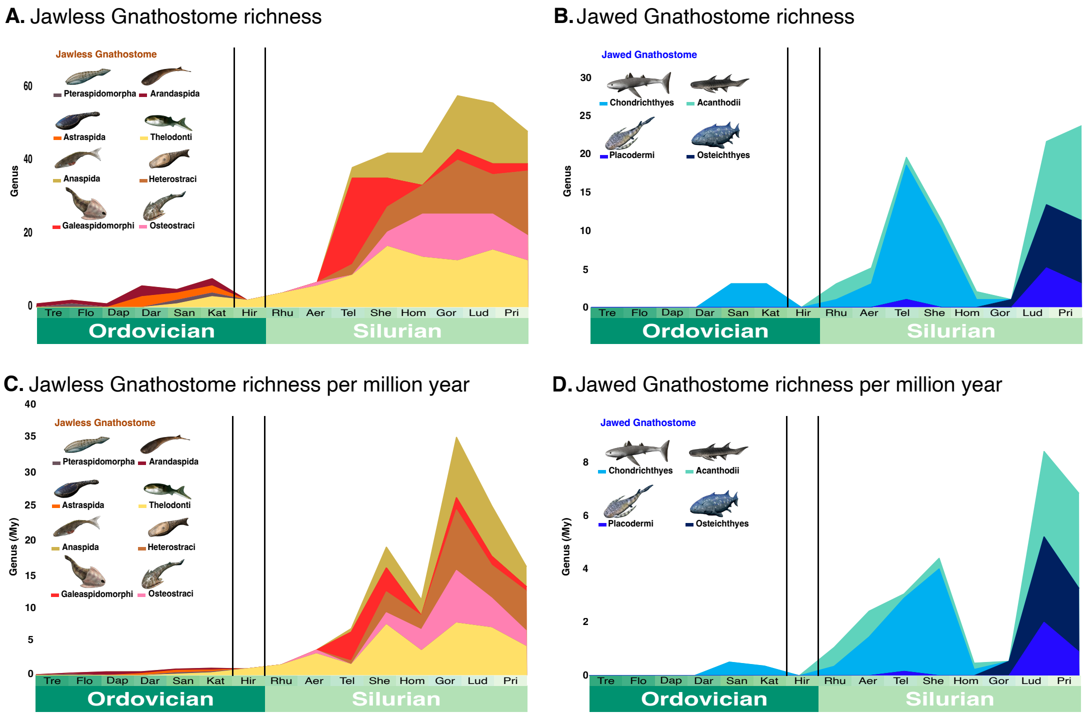


**Fig. S3. Genus-level richness diversity curves for Ordovician-Silurian jawed and jawless gnathostome groups. (A) and (C)** Jawless gnathostome groups (N = 318). **(B) and (D)** Jawed-gnathostome groups (N = 90). **(A) and (B)** Raw number of genera in each stage (Data range:1–59; 0–23). **(C) and (D)** Number of genera per million years in each stage (Data range:0.10–34.71; 0–9.13).

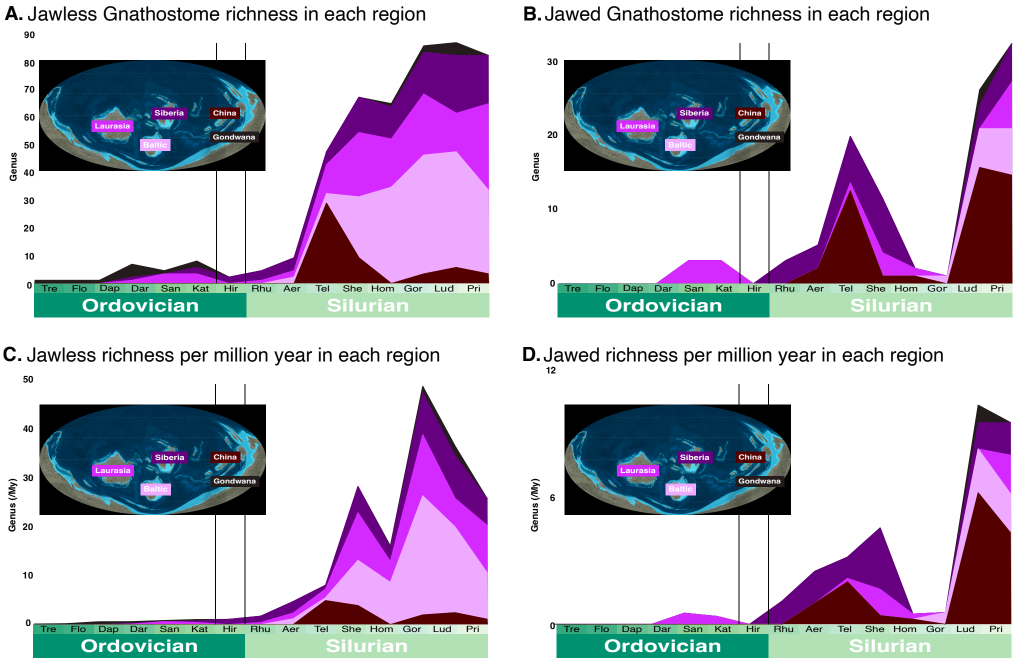


**Figure S4. Diversity curves for jawed and jawless gnathostomes richness in each region from Ordovician to Silurian at genus level. (A) to (D)** Genus-level richness per stage divided by region. **(A)** Raw genus-level richness per stage for jawless gnathostomes in each region (N = 387; Data range:1–75). **(B)** Raw genus-level richness per stage for jawed gnathostomes in each region (N = 97; Data range: 0–31). **(C)** Stage-level jawless gnathostome genus richness per million years in each region **(D)** Stage-level jawless gnathostome genus richness per million years in each region. Paleomaps from Deep Time Maps and used with permission (*54*).

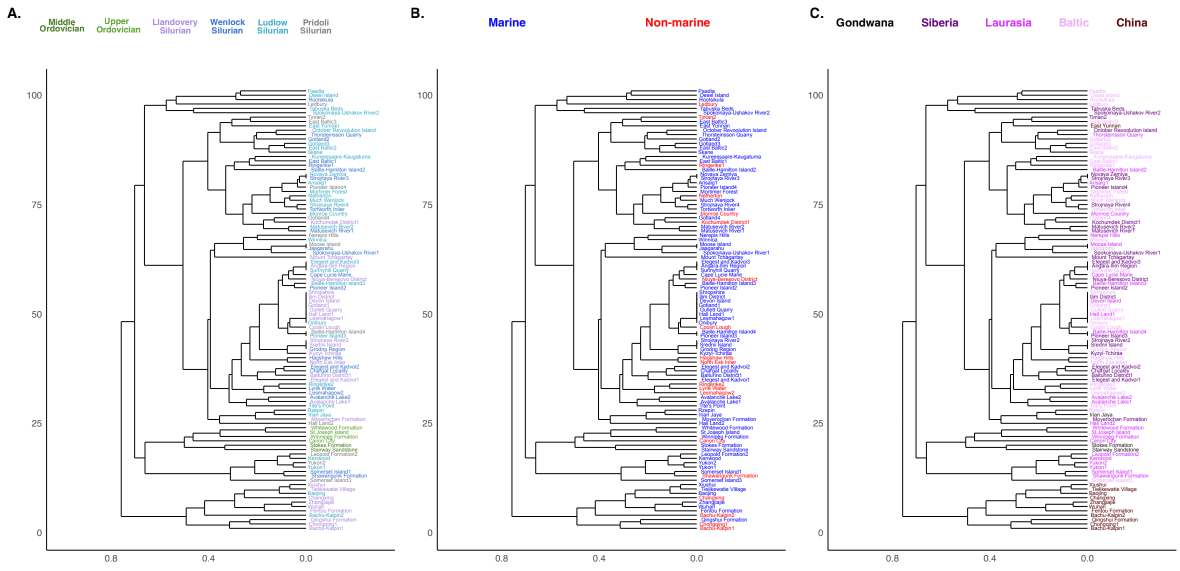


**Figure S5. Cluster dendrogram of Ordovician-Silurian assemblages based on species-level diversity within groups.** Cluster dendrogram of 101 gnathostome assemblages based on Bray-Curtis distance and UPGMA (Cophenetic correlation: 0.85). **(A):** Assemblages colored using the same scheme as in Fig. 2B–C for clarity based on series **(B)** Assemblages colored by environment (marine vs. non-marine). **(C)** Assemblages colored by region.

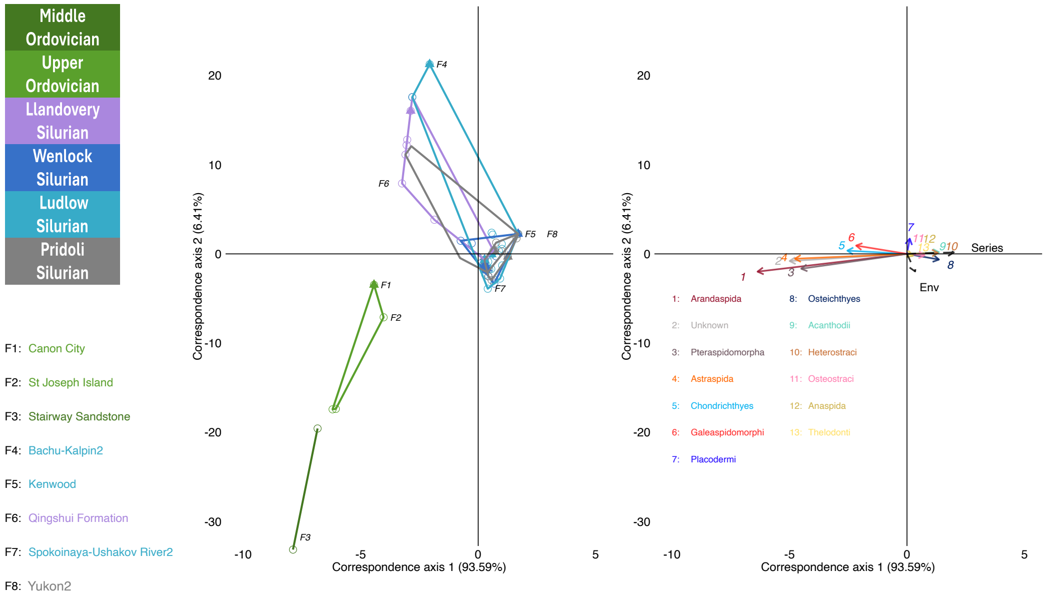


**Figure S6. Canonical Correspondence Analysis plots (CCA) for Ordovician-Silurian gnathostome assemblages.** This graph is the same as in Fig. 2B, but showed the position of all gnathostome groups in right panel.

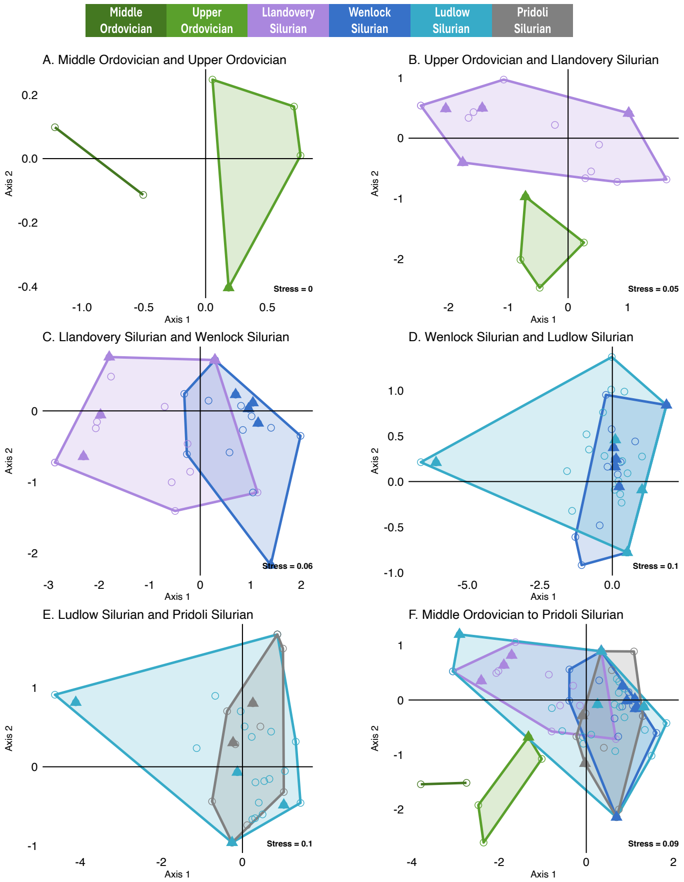


**Figure S7. Non-parametric multidimensional scaling (NMDS) plots for gnathostome assemblages (raw species richness, Bray-Curtis distance). (A)** Middle Ordovician (N = 2) and Upper Ordovician (N = 4). **(B)** Upper Ordovician (N = 4) and Llandovery (N = 29) **(C)** Llandovery (N = 29) and Wenlock (N = 22) **(D)** Wenlock (N = 22) and Ludlow (N = 31) **(E)** Ludlow (N = 31) and Pridoli (N = 11) **(F)** All Ordovician-Silurian assemblages (N = 101).

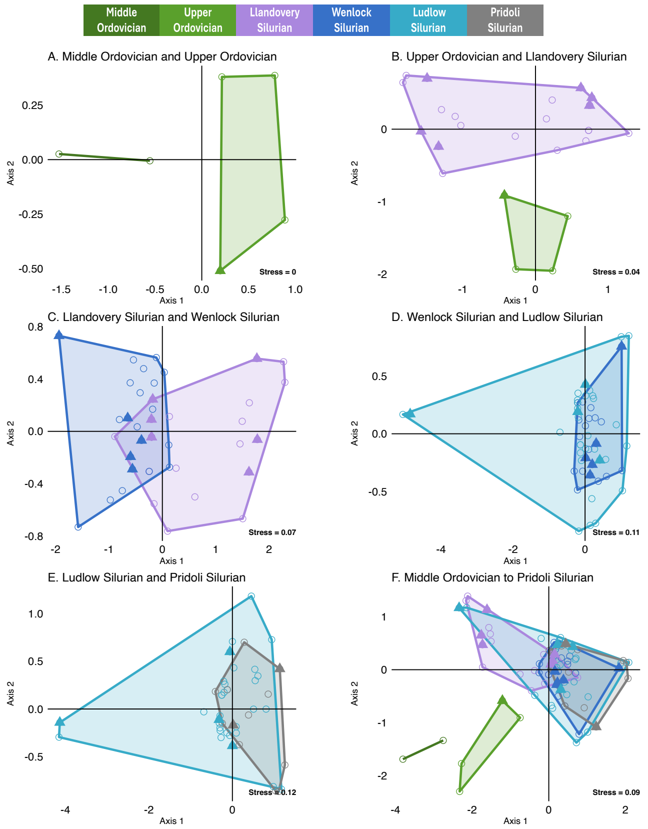


**Figure S8. NMDS for Ordovician to Silurian gnathostome assemblages (group presence-absence, Kulczynski dissimilarity) (A)** Middle Ordovician (N = 2) and Upper Ordovician (N = 4). **(B)** Upper Ordovician (N = 4) and Llandovery (N = 29) **(C)** Llandovery (N = 29) and Wenlock (N = 22) **(D)** Wenlock (N = 22) and Ludlow (N = 31) **(E)** Ludlow (N = 31) and Pridoli (N=11) **(F)** All Ordovician-Silurian assemblages (N = 101).

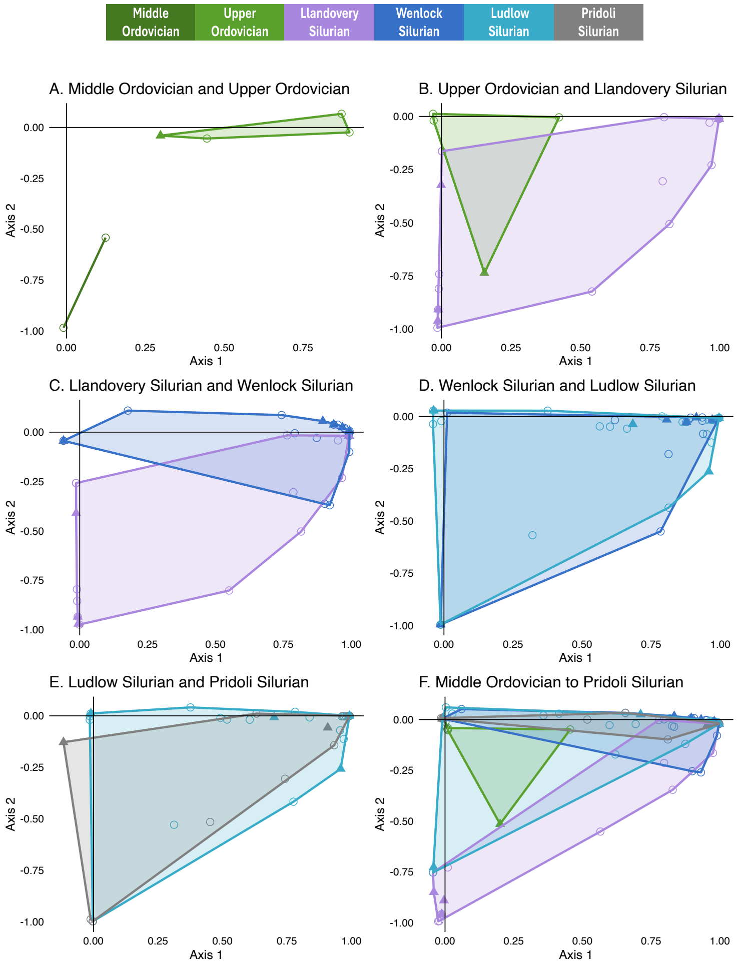


**Figure S9. Factor Analysis for Ordovician to Silurian in raw diversity.** FA for sites in each stage. **(A)** Middle Ordovician (N = 2) and Upper Ordovician (N = 4). **(B)** Upper Ordovician (N = 4) and Llandovery (N = 29) **(C)** Llandovery (N = 29) and Wenlock (N = 22) **(D)** Wenlock (N = 22) and Ludlow (N = 31) **(E)** Ludlow (N = 31) and Pridoli (N=11) **(F)** All Ordovician-Silurian assemblages (N = 101).

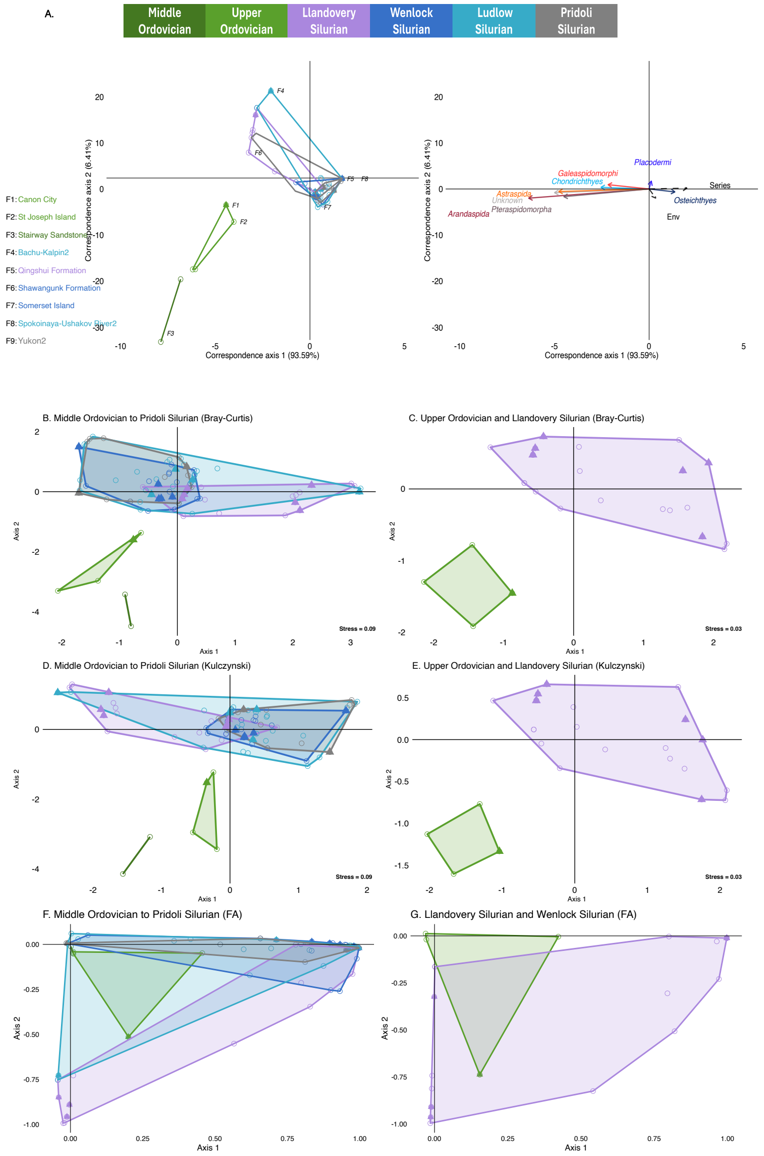


**Figure S10. Ordination and FA plots of assemblages when Ordovician Chondrichthyes reassigned as "Unknown." (reanalysis 1) (A)** CCA of raw species occurrences for 101 localities (N = 1,076). Non-marine assemblages were marked with filled triangles. **Left:** site ordination plot, with 9 representative faunas (assemblages) annotated in the caption, including those showing the maximum and minimum scores along each axis. Sites *F5* and *F8* share the same score and thus overlap at the same coordinate. **Right:** taxon ordination plot (N = 13), showing the top 7 Paleozoic fish groups with the highest contributions, displayed as biplot arrows. **(B) and (C)** NMDS analysis based on raw diversity and Bray-Curtis distances. **(B)** Plot for Ordovician to Silurian. **(C)** Plot for Upper-Ordovician to Llandovery Silurian. **(D) and (E)** NMDS analysis based on presence-absence for groups and Kulczynski dissimilarity. **(D)** plot for Ordovician to Silurian. **(E)** plot for Upper-Ordovician to Llandovery Silurian**. (F) and (G)** Factor analysis (FA) for assemblages. **(F)** plot for Ordovician to Silurian; **(G)** plot for Upper-Ordovician to Llandovery Silurian.

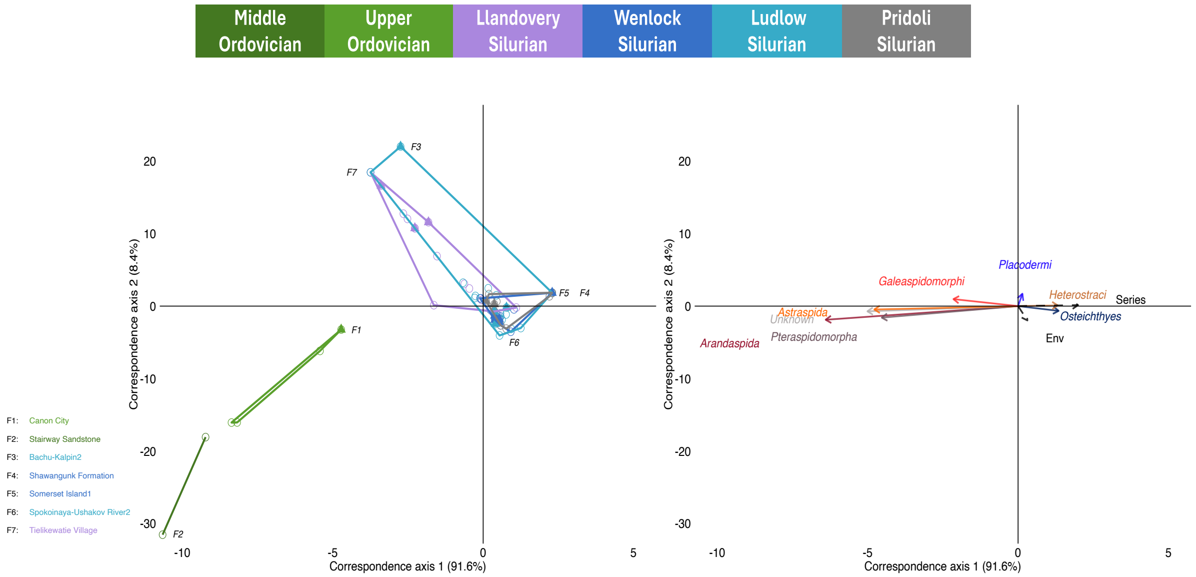


**Figure S11. CCA of gnathostome assemblages with all Acanthodii grouped as Chondrichthyes (reanalysis 2).** Raw species occurrences for 101 localities were used (N = 1,076). Non-marine sites were marked with filled triangles. **Left:** Assemblage ordination plot, with 9 representative faunas annotated in the caption, including those showing the maximum and minimum scores along each axis. Sites *F4* and *F5* share the same score and thus overlap at the same coordinate. **Right:** Taxon ordination plot (N = 12), showing the top 7 Paleozoic fish groups with the highest contributions, displayed as biplot arrows.

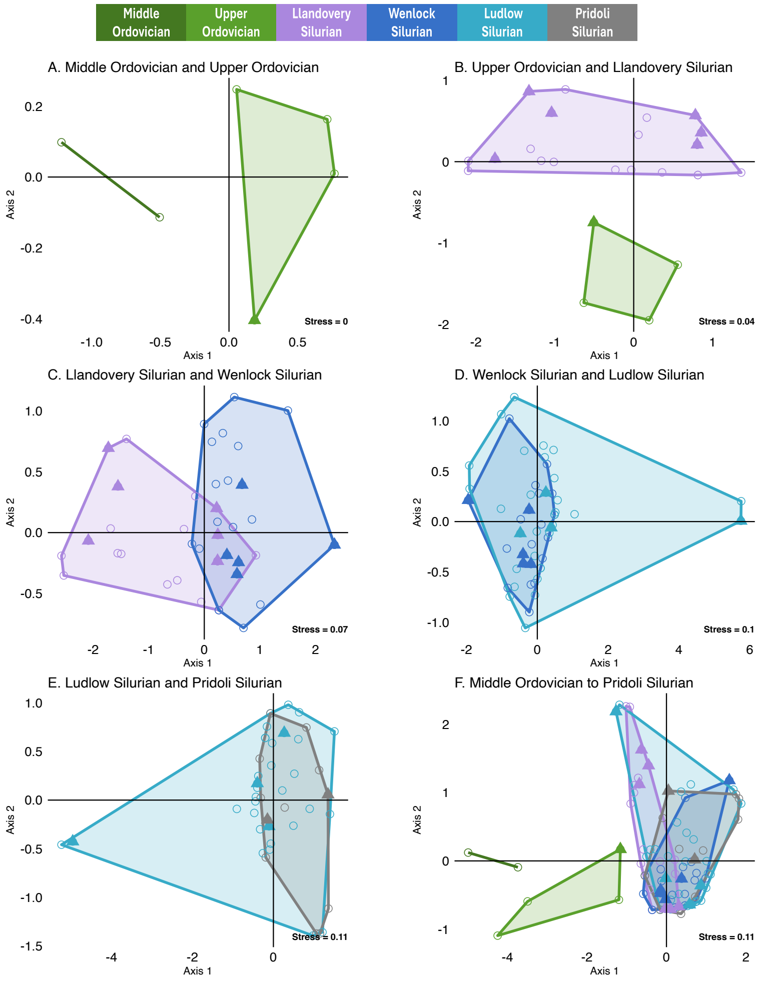


**Figure S12. NMDS of gnathostome assemblages with all Acanthodii reassigned as Chondrichthyes (reanalysis 2) (raw species diversity, Bray-Curtis distance)** NMDS for sites in each stage. **(A)** Middle Ordovician (N = 2) and Upper Ordovician (N = 4). **(B)** Upper Ordovician (N = 4) and Llandovery (N = 29) **(C)** Llandovery (N = 29) and Wenlock (N = 22) **(D)** Wenlock (N = 22) and Ludlow (N = 31) **(E)** Ludlow (N = 31) and Pridoli (N=11) **(F)** All Ordovician-Silurian assemblages (N = 101).

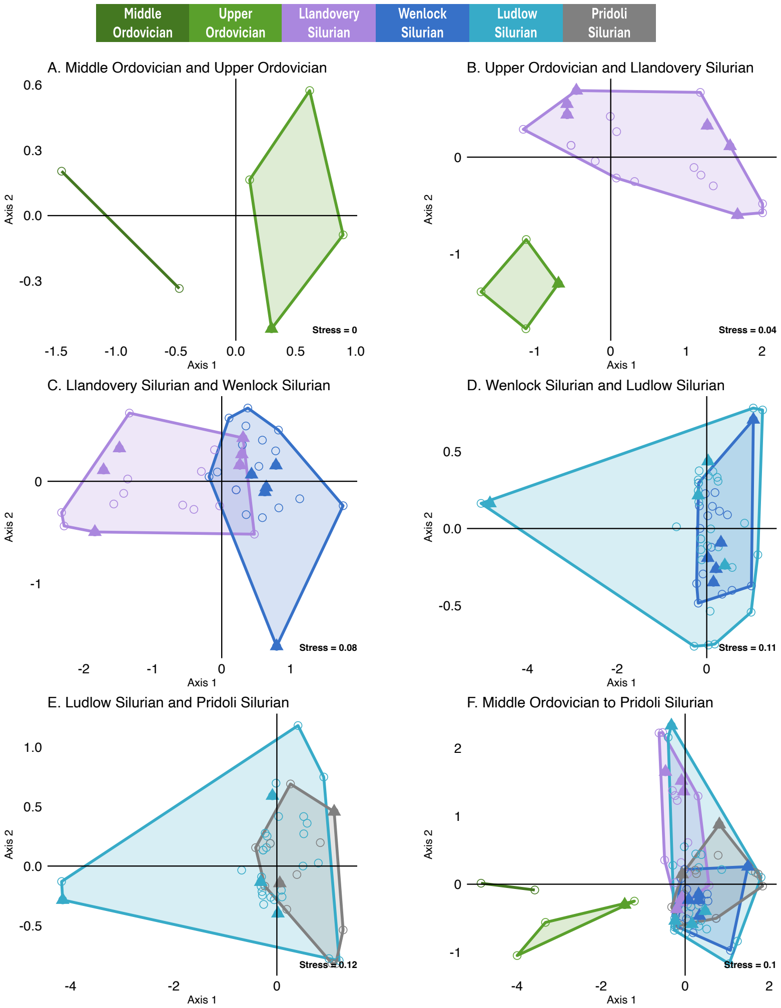


**Figure S13. NMDS of gnathostome assemblages with all Acanthodii reassigned as Chondrichthyes (reanalysis 2) (group presence-absence, Kulczynski dissimilarity).** NMDS for assemblages in each stage. **(A)** Middle Ordovician (N = 2) and Upper Ordovician (N = 4). **(B)** Upper Ordovician (N = 4) and Llandovery (N = 29) **(C)** Llandovery (N = 29) and Wenlock (N = 22) **(D)** Wenlock (N = 22) and Ludlow (N = 31) **(E)** Ludlow (N = 31) and Pridoli (N=11) **(F)** All Ordovician-Silurian assemblages (N = 101).

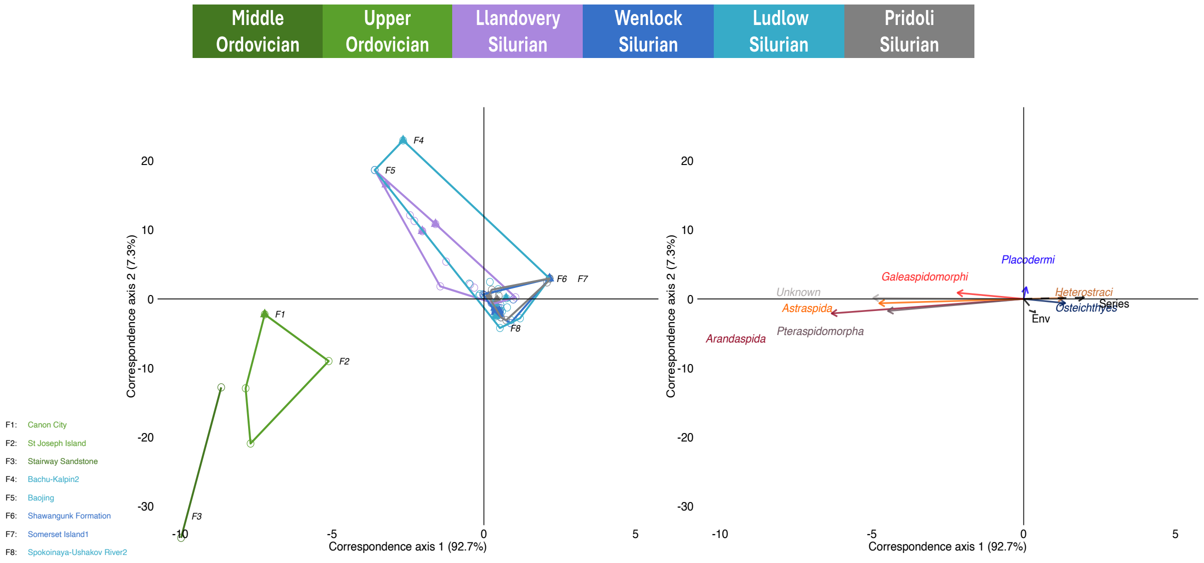


**Figure S14. CCA of gnathostome assemblages with all Silurian Acanthodii treated as Chondrichthyes and all Ordovician Chondrichthyes treated as "Unknown" (reanalysis 3).** Raw species occurrences for 101 localities were used (N = 1,076). Non-marine sites were marked with filled triangles. **Left:** Assemblage ordination plot, with 9 representative faunas annotated in the caption, including those showing the maximum and minimum scores along each axis. Sites *F4* and *F5* share the same score and thus overlap at the same coordinate. **Right:** Taxon ordination plot (N = 12), showing the top 7 Paleozoic fish groups with the highest contributions, displayed as biplot arrows.

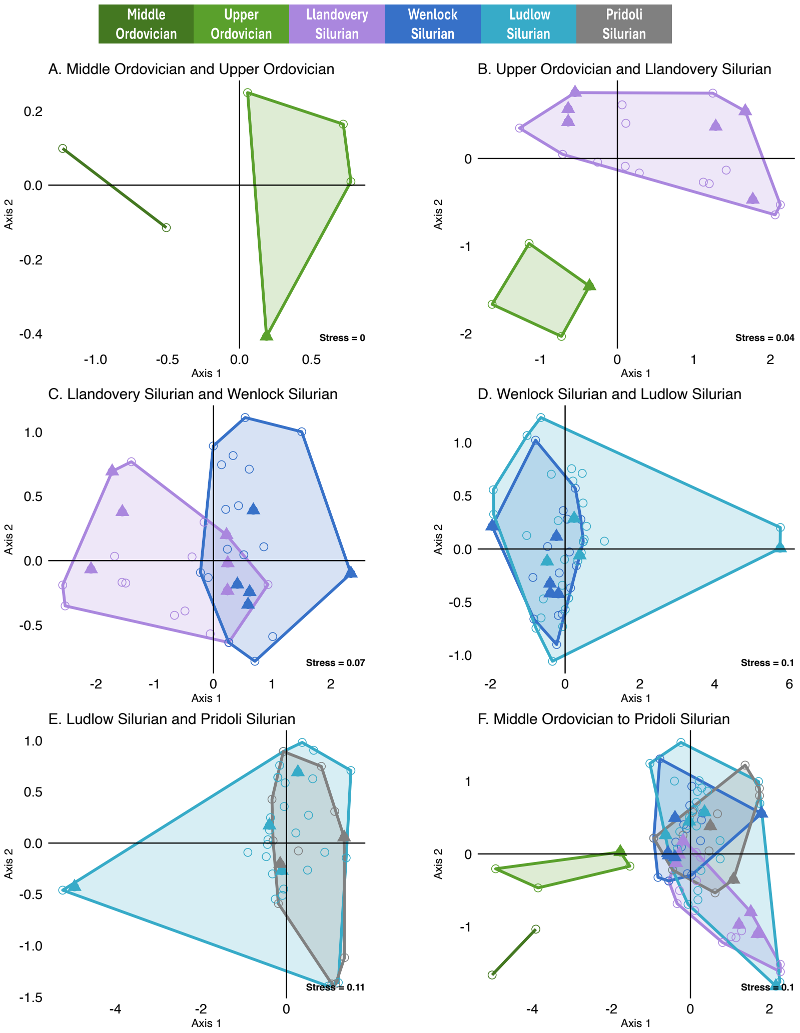


**Figure S15. NMDS of gnathostome assemblages with all Silurian Acanthodii treated as Chondrichthyes and all Ordovician Chondrichthyes treated as "Unknown" (reanalysis 3) (raw species richness, Bray-Curtis distance).** NMDS for sites in each stage. **(A)** Middle Ordovician (N = 2) and Upper Ordovician (N = 4). **(B)** Upper Ordovician (N = 4) and Llandovery (N = 29) **(C)** Llandovery (N = 29) and Wenlock (N = 22) **(D)** Wenlock (N = 22) and Ludlow (N = 31) **(E)** Ludlow (N = 31) and Pridoli (N=11) **(F)** All Ordovician-Silurian assemblages (N = 101).

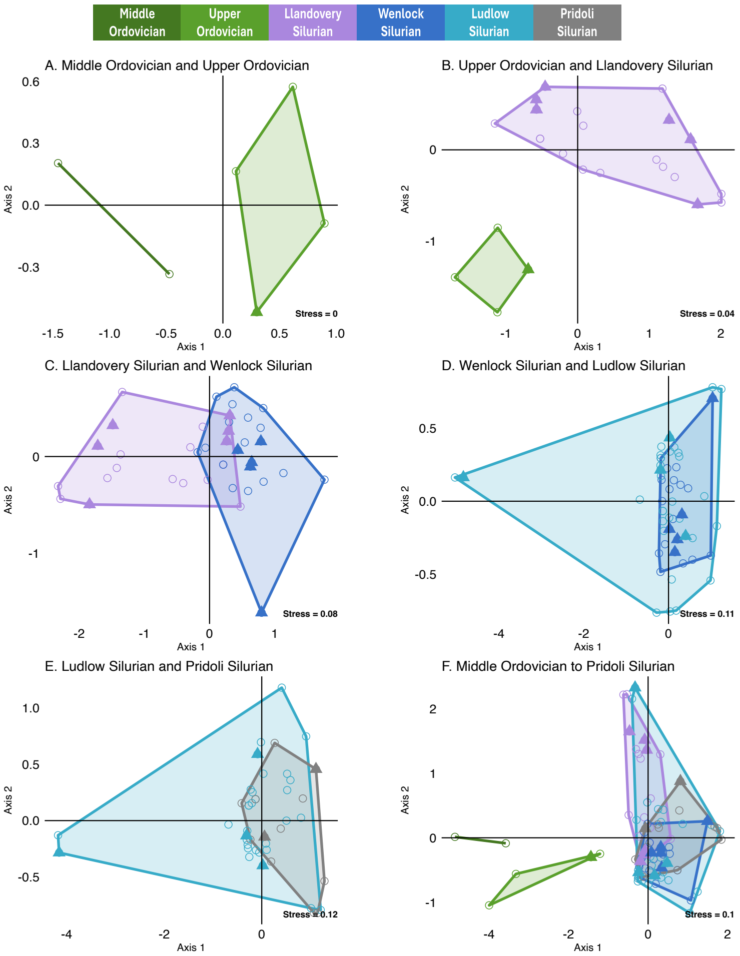


**Figure S16. NMDS of gnathostome assemblages with all Silurian Acanthodii treated as Chondrichthyes and all Ordovician Chondrichthyes treated as "Unknown" (reanalysis 3) (group presence-absence, Kulczynski dissimilarity).** NMDS for sites in each stage. **(A)** Middle Ordovician (N = 2) and Upper Ordovician (N = 4). **(B)** Upper Ordovician (N = 4) and Llandovery (N = 29) **(C)** Llandovery (N = 29) and Wenlock (N = 22) **(D)** Wenlock (N = 22) and Ludlow (N = 31) **(E)** Ludlow (N = 31) and Pridoli (N=11) **(F)** All Ordovician-Silurian assemblages (N = 101).

**Table S1. Occurrences of 13 Paleozoic gnathostome groups at the species level. (A)** The raw data included 59 species occurrences from the Ordovician. **(B)** The raw data included 1,098 species occurrences from the Silurian. These data were used for Fig. 1. The values marked with an asterisk (*) indicate that the occurrences counted twice due to the presence of the same fauna across two different stages.

A.

| Class | Tremadocian | Floian | Dapingian | Darriwilian | Sandbian | Katian | Hirnantian |
|---|---|---|---|---|---|---|---|
| Pteraspidomorpha | 0 | 1 | 0 | 0 | 2 | 1 | 0 |
| Arandaspida | 1 | *2 | 2 | *11 | 3 | 1 | 0 |
| Astraspida | 0 | 0 | 0 | 3 | 9 | *13 | 0 |
| Heterostraci | 0 | 0 | 0 | 0 | 0 | 0 | 0 |
| Galeaspidomorphi | 0 | 0 | 0 | 0 | 0 | 0 | 0 |
| Anaspida | 0 | 0 | 0 | 0 | 0 | 0 | 0 |
| Thelodonti | 0 | 0 | 0 | 0 | 2 | *5 | *3 |
| Osteostraci | 0 | 0 | 0 | 0 | 0 | 0 | 0 |
| Chondricthyes | 0 | 0 | 0 | 0 | 4 | *4 | 0 |
| Acanthodii | 0 | 0 | 0 | 0 | 0 | 0 | 0 |
| Placodermi | 0 | 0 | 0 | 0 | 0 | 0 | 0 |
| Osteichthyes | 0 | 0 | 0 | 0 | 0 | 0 | 0 |
| Unknown | 0 | 0 | 0 | 3 | 5 | *2 | 1 |
| Total | 1 | *3 | 2 | *17 | 25 | *26 | *4 |

B.

| Class | Rhuddanian | Aeronian | Telychian | Sheinwoodian | Homerian | Gorstian | Ludfordian | Pridoli |
|---|---|---|---|---|---|---|---|---|
| Pteraspidomorpha | 0 | 0 | 0 | 0 | 0 | 0 | 0 | 0 |
| Arandaspida | 0 | 0 | 0 | 0 | 0 | 0 | 0 | 0 |
| Astraspida | 0 | 0 | 0 | 0 | 0 | 0 | 0 | 0 |
| Heterostraci | 0 | 0 | 3 | *11 | *14 | *33 | *30 | *38 |
| Galeaspidomorphi | 0 | 0 | 39 | *11 | 0 | 3 | 5 | *2 |
| Anaspida | 0 | 0 | 4 | *35 | *43 | *40 | *44 | *20 |
| Thelodonti | 9 | *49 | *95 | *156 | *119 | *190 | *271 | *176 |
| Osteostraci | 0 | 1 | 0 | 8 | *18 | *43 | *39 | *17 |
| Chondricthyes | 1 | *3 | 47 | *11 | 1 | 0 | 1 | *1 |
| Acanthodii | 3 | *3 | *3 | 1 | 1 | 1 | *23 | *43 |
| Placodermi | 0 | 0 | 1 | 0 | 0 | 1 | 5 | *3 |
| Osteichthyes | 0 | 0 | 0 | 0 | 0 | 3 | *11 | *10 |
| Unknown | *2 | *1 | 0 | 0 | 0 | 0 | 0 | 0 |
| Total | *15 | *57 | *192 | *233 | *196 | *314 | *429 | *310 |

**Table S2. Species-level occurrences for 13 Paleozoic gnathostome groups in each region and series.** These data were used in Fig. 3. **(A)** Lower Ordovician (N = 3). **(B)** Middle Ordovician (N = 17). **(C)** Upper Ordovician (N = 39). **(D)** Llandovery Silurian (N = 218). **(E)** Wenlock Silurian (N = 274). **(F)** Ludlow Silurian (N = 495). **(G)** Pridoli Silurian (N = 310).

A.

| | **Gondwana** | **Siberia** | **Laurasia** | **Baltic** | **China** |
|---|---|---|---|---|---|
| **Pteraspidomorpha** | 1 | 0 | 0 | 0 | 0 |
| **Arandaspida** | 2 | 0 | 0 | 0 | 0 |
| **Astraspida** | 0 | 0 | 0 | 0 | 0 |
| **Heterostraci** | 0 | 0 | 0 | 0 | 0 |
| **Galeaspidomorphi** | 0 | 0 | 0 | 0 | 0 |
| **Anaspida** | 0 | 0 | 0 | 0 | 0 |
| **Thelodonti** | 0 | 0 | 0 | 0 | 0 |
| **Osteostraci** | 0 | 0 | 0 | 0 | 0 |
| **Chondrichthyes** | 0 | 0 | 0 | 0 | 0 |
| **Acanthodii** | 0 | 0 | 0 | 0 | 0 |
| **Placodermi** | 0 | 0 | 0 | 0 | 0 |
| **Osteichthyes** | 0 | 0 | 0 | 0 | 0 |
| **Unknown** | 0 | 0 | 0 | 0 | 0 |

B.

| | **Gondwana** | **Siberia** | **Laurasia** | **Baltic** | **China** |
|---|---|---|---|---|---|
| **Pteraspidomorpha** | 0 | 0 | 0 | 0 | 0 |
| **Arandaspida** | 11 | 0 | 0 | 0 | 0 |
| **Astraspida** | 1 | 1 | 1 | 0 | 0 |
| **Heterostraci** | 0 | 0 | 0 | 0 | 0 |
| **Galeaspidomorphi** | 0 | 0 | 0 | 0 | 0 |
| **Anaspida** | 0 | 0 | 0 | 0 | 0 |
| **Thelodonti** | 0 | 0 | 0 | 0 | 0 |
| **Osteostraci** | 0 | 0 | 0 | 0 | 0 |
| **Chondrichthyes** | 0 | 0 | 0 | 0 | 0 |
| **Acanthodii** | 0 | 0 | 0 | 0 | 0 |
| **Placodermi** | 0 | 0 | 0 | 0 | 0 |
| **Osteichthyes** | 0 | 0 | 0 | 0 | 0 |
| **Unknown** | 3 | 0 | 0 | 0 | 0 |

C.

| | Gondwana | Siberia | Laurasia | Baltic | China |
|---|---|---|---|---|---|
| **Pteraspidomorpha** | 0 | 0 | 3 | 0 | 0 |
| **Arandaspida** | 4 | 0 | 0 | 0 | 0 |
| **Astraspida** | 1 | 0 | 15 | 0 | 0 |
| **Heterostraci** | 0 | 0 | 0 | 0 | 0 |
| **Galeaspidomorphi** | 0 | 0 | 0 | 0 | 0 |
| **Anaspida** | 0 | 0 | 0 | 0 | 0 |
| **Thelodonti** | 0 | 4 | 2 | 0 | 0 |
| **Osteostraci** | 0 | 0 | 0 | 0 | 0 |
| **Chondrichthyes** | 0 | 0 | 4 | 0 | 0 |
| **Acanthodii** | 0 | 0 | 0 | 0 | 0 |
| **Placodermi** | 0 | 0 | 0 | 0 | 0 |
| **Osteichthyes** | 0 | 0 | 0 | 0 | 0 |
| **Unknown** | 0 | 1 | 5 | 0 | 0 |

D.

| | Gondwana | Siberia | Laurasia | Baltic | China |
|---|---|---|---|---|---|
| **Pteraspidomorpha** | 0 | 0 | 0 | 0 | 0 |
| **Arandaspida** | 0 | 0 | 0 | 0 | 0 |
| **Astraspida** | 0 | 0 | 0 | 0 | 0 |
| **Heterostraci** | 0 | 0 | 3 | 0 | 0 |
| **Galeaspidomorphi** | 0 | 0 | 0 | 0 | 39 |
| **Anaspida** | 0 | 0 | 2 | 2 | 0 |
| **Thelodonti** | 0 | 57 | 22 | 33 | 0 |
| **Osteostraci** | 0 | 0 | 0 | 1 | 0 |
| **Chondrichthyes** | 0 | 9 | 0 | 0 | 41 |
| **Acanthodii** | 0 | 6 | 0 | 0 | 0 |
| **Placodermi** | 0 | 0 | 0 | 0 | 1 |
| **Osteichthyes** | 0 | 0 | 0 | 0 | 0 |
| **Unknown** | 0 | 2 | 0 | 0 | 0 |

E.

| | Gondwana | Siberia | Laurasia | Baltic | China |
|---|---|---|---|---|---|
| **Pteraspidomorpha** | 0 | 0 | 0 | 0 | 0 |
| **Arandaspida** | 0 | 0 | 0 | 0 | 0 |
| **Astraspida** | 0 | 0 | 0 | 0 | 0 |
| **Heterostraci** | 0 | 0 | 17 | 1 | 0 |
| **Galeaspidomorphi** | 0 | 0 | 0 | 0 | 11 |
| **Anaspida** | 0 | 5 | 12 | 28 | 0 |
| **Thelodonti** | 1 | 48 | 49 | 70 | 0 |
| **Osteostraci** | 0 | 1 | 0 | 18 | 0 |
| **Chondrichthyes** | 0 | 7 | 3 | 0 | 1 |
| **Acanthodii** | 0 | 1 | 0 | 0 | 1 |
| **Placodermi** | 0 | 0 | 0 | 0 | 0 |
| **Osteichthyes** | 0 | 0 | 0 | 0 | 0 |
| **Unknown** | 0 | 0 | 0 | 0 | 0 |

F.

| | Gondwana | Siberia | Laurasia | Baltic | China |
|---|---|---|---|---|---|
| **Pteraspidomorpha** | 0 | 0 | 0 | 0 | 0 |
| **Arandaspida** | 0 | 0 | 0 | 0 | 0 |
| **Astraspida** | 0 | 0 | 0 | 0 | 0 |
| **Heterostraci** | 0 | 3 | 27 | 10 | 0 |
| **Galeaspidomorphi** | 0 | 0 | 0 | 0 | 8 |
| **Anaspida** | 0 | 16 | 2 | 37 | 0 |
| **Thelodonti** | 5 | 70 | 37 | 174 | 16 |
| **Osteostraci** | 0 | 1 | 0 | 48 | 0 |
| **Chondrichthyes** | 1 | 0 | 0 | 0 | 0 |
| **Acanthodii** | 5 | 2 | 0 | 10 | 6 |
| **Placodermi** | 0 | 0 | 0 | 0 | 6 |
| **Osteichthyes** | 0 | 1 | 0 | 4 | 6 |
| **Unknown** | 0 | 0 | 0 | 0 | 0 |

G.

| | Gondwana | Siberia | Laurasia | Baltic | China |
|---|---|---|---|---|---|
| **Pteraspidomorpha** | 0 | 0 | 0 | 0 | 0 |
| **Arandaspida** | 0 | 0 | 0 | 0 | 0 |
| **Astraspida** | 0 | 0 | 0 | 0 | 0 |
| **Heterostraci** | 0 | 4 | 17 | 17 | 0 |
| **Galeaspidomorphi** | 0 | 0 | 0 | 0 | 2 |
| **Anaspida** | 0 | 5 | 3 | 12 | 0 |
| **Thelodonti** | 0 | 46 | 16 | 98 | 16 |
| **Osteostraci** | 0 | 1 | 2 | 14 | 0 |
| **Chondrichthyes** | 0 | 0 | 1 | 0 | 0 |
| **Acanthodii** | 0 | 8 | 8 | 21 | 6 |
| **Placodermi** | 0 | 0 | 0 | 0 | 3 |
| **Osteichthyes** | 0 | 1 | 0 | 3 | 6 |
| **Unknown** | 0 | 0 | 0 | 0 | 0 |

**Table S3. Genus-level richness per stage for 13 Paleozoic gnathostome groups, total group gnathostomes and conodonts.** These data were used in Fig. 1. Total group gnathostome richness was counted using 12 Paleozoic groups excluding the "Unknown" category. The values marked with an asterisk (*) indicate that the occurrences counted twice due to the presence of the same fauna across two different stages. **(A)** Ordovician (gnathostomes N = 31, conodonts N = 363) **(B)** Silurian (gnathostomes N = 386, conodonts N = 117) **(C)** Ordovician genus richness divided by million years for each stage. **(D)** Silurian genus richness divided by million years for each stage.

A.

| Class | Tremadocian | Floian | Dapingian | Darriwilian | Sandbian | Katian | Hirnantian |
|---|---|---|---|---|---|---|---|
| Pteraspidomorpha | 0 | 1 | 0 | 0 | 1 | 1 | 0 |
| Arandaspida | 1 | 1 | 1 | 3 | 1 | 2 | 0 |
| Astraspida | 0 | 0 | 0 | 3 | 2 | 2 | 0 |
| Heterostraci | 0 | 0 | 0 | 0 | 0 | 0 | 0 |
| Galeaspidomorphi | 0 | 0 | 0 | 0 | 0 | 0 | 0 |
| Anaspida | 0 | 0 | 0 | 0 | 0 | 0 | 0 |
| Thelodonti | 0 | 0 | 0 | 0 | 1 | 3 | *2 |
| Osteostraci | 0 | 0 | 0 | 0 | 0 | 0 | 0 |
| Chondricthyes | 0 | 0 | 0 | 0 | 3 | *3 | 0 |
| Acanthodii | 0 | 0 | 0 | 0 | 0 | 0 | 0 |
| Placodermi | 0 | 0 | 0 | 0 | 0 | 0 | 0 |
| Osteichthyes | 0 | 0 | 0 | 0 | 0 | 0 | 0 |
| Unknown | 0 | 0 | 0 | 2 | 2 | *2 | 1 |
| Gnathostome | 1 | 2 | 1 | 6 | 8 | *11 | *2 |
| Conodonts | 51 | 69 | 23 | 77 | 64 | 61 | 18 |

B.

| Class | Rhuddanian | Aeronian | Telychian | Sheinwoodian | Homerian | Gorstian | Ludfordian | Pridoli |
|---|---|---|---|---|---|---|---|---|
| Pteraspidomorpha | 0 | 0 | 0 | 0 | 0 | 0 | 0 | 0 |
| Arandaspida | 0 | 0 | 0 | 0 | 0 | 0 | 0 | 0 |
| Astraspida | 0 | 0 | 0 | 0 | 0 | 0 | 0 | 0 |
| Heterostraci | 0 | 0 | 3 | *7 | *8 | *15 | *11 | 18 |
| Galeaspidomorphi | 0 | 0 | 24 | *8 | 0 | 3 | 3 | *2 |
| Anaspida | 0 | 0 | 3 | 7 | *9 | 15 | *17 | *9 |
| Thelodonti | 4 | 6 | *9 | 17 | *14 | 13 | *16 | *13 |
| Osteostraci | 0 | 1 | 0 | 4 | 12 | *13 | *10 | *7 |
| Chondricthyes | 1 | *3 | 17 | *10 | *1 | 0 | 0 | 0 |
| Acanthodii | 2 | *2 | 1 | 1 | 1 | 0 | 8 | *12 |
| Placodermi | 0 | 0 | 1 | 0 | 0 | 0 | 5 | 3 |
| Osteichthyes | 0 | 0 | 0 | 0 | 0 | 1 | 8 | *8 |
| Unknown | 1 | *1 | 0 | 0 | 0 | 0 | 0 | 0 |
| Gnathostome | 7 | *12 | *58 | *54 | *45 | *60 | *78 | *72 |
| Conodonts | 13 | 9 | 23 | 18 | 10 | 10 | 15 | 19 |

C.

| Class | Tremadocian | Floian | Dapingian | Darriwilian | Sandbian | Katian | Hirnantian |
|---|---|---|---|---|---|---|---|
| **Pteraspidomorpha** | 0.00 | 0.17 | 0.00 | 0.00 | 0.19 | 0.13 | 0.00 |
| **Arandaspida** | 0.10 | 0.17 | 0.53 | 0.27 | 0.19 | 0.26 | 0.00 |
| **Astraspida** | 0.00 | 0.00 | 0.00 | 0.27 | 0.37 | 0.26 | 0.00 |
| **Heterostraci** | 0.00 | 0.00 | 0.00 | 0.00 | 0.00 | 0.00 | 0.00 |
| **Galeaspidomorphi** | 0.00 | 0.00 | 0.00 | 0.00 | 0.00 | 0.00 | 0.00 |
| **Anaspida** | 0.00 | 0.00 | 0.00 | 0.00 | 0.00 | 0.00 | 0.00 |
| **Thelodonti** | 0.00 | 0.00 | 0.00 | 0.00 | 0.19 | 0.39 | *0.95 |
| **Osteostraci** | 0.00 | 0.00 | 0.00 | 0.00 | 0.00 | 0.00 | 0.00 |
| **Chondricthyes** | 0.00 | 0.00 | 0.00 | 0.00 | 0.56 | *0.39 | 0.00 |
| **Acanthodii** | 0.00 | 0.00 | 0.00 | 0.00 | 0.00 | 0.00 | 0.00 |
| **Placodermi** | 0.00 | 0.00 | 0.00 | 0.00 | 0.00 | 0.00 | 0.00 |
| **Osteichthyes** | 0.00 | 0.00 | 0.00 | 0.00 | 0.00 | 0.00 | 0.00 |
| **Unknown** | 0.00 | 0.00 | 0.00 | 0.18 | 0.37 | *0.26 | 0.48 |
| **Gnathostome** | 0.10 | 0.34 | 0.53 | 0.54 | 1.48 | *1.45 | *0.95 |
| **Conodonts** | 5.20 | 11.90 | 12.11 | 6.88 | 11.85 | 8.03 | 8.57 |

D.

| Class | Rhuddanian | Aeronian | Telychian | Sheinwoodian | Homerian | Gorstian | Ludfordian | Pridoli |
|---|---|---|---|---|---|---|---|---|
| **Pteraspidomorpha** | 0.00 | 0.00 | 0.00 | 0.00 | 0.00 | 0.00 | 0.00 | 0.00 |
| **Arandaspida** | 0.00 | 0.00 | 0.00 | 0.00 | 0.00 | 0.00 | 0.00 | 0.00 |
| **Astraspida** | 0.00 | 0.00 | 0.00 | 0.00 | 0.00 | 0.00 | 0.00 | 0.00 |
| **Heterostraci** | 0.00 | 0.00 | 0.53 | *3.04 | *2.05 | *8.82 | *4.78 | 5.81 |
| **Galeaspidomorphi** | 0.00 | 0.00 | 4.21 | *3.48 | 0.00 | 1.76 | 1.30 | *0.65 |
| **Anaspida** | 0.00 | 0.00 | 0.53 | 3.04 | *2.31 | 8.82 | *7.39 | *2.90 |
| **Thelodonti** | 1.54 | 3.16 | *1.58 | 7.39 | *3.59 | 7.65 | *6.96 | *4.19 |
| **Osteostraci** | 0.00 | 0.53 | 0.00 | 1.74 | 3.08 | *7.65 | *4.35 | *2.26 |
| **Chondricthyes** | 0.38 | *1.58 | 2.98 | *4.35 | *0.26 | 0.00 | 0.00 | 0.00 |
| **Acanthodii** | 0.77 | *1.05 | 0.18 | 0.43 | 0.26 | 0.00 | 3.48 | *3.87 |
| **Placodermi** | 0.00 | 0.00 | 0.18 | 0.00 | 0.00 | 0.00 | 2.17 | 0.97 |
| **Osteichthyes** | 0.00 | 0.00 | 0.00 | 0.00 | 0.00 | 0.59 | 3.48 | *2.58 |
| **Unknown** | 0.38 | *0.53 | 0.00 | 0.00 | 0.00 | 0.00 | 0.00 | 0.00 |
| **Gnathostome** | 2.69 | *6.32 | *10.18 | *23.48 | *11.54 | *35.29 | *33.91 | *23.23 |
| **Conodonts** | 5.00 | 4.74 | 4.04 | 7.83 | 2.56 | 5.88 | 6.52 | 6.13 |

**Table S4. The number of assemblages (faunas) per stage in 5 regions. (A)** Ordovician (N = 37) **(B)** Silurian (N =202). The values marked with an asterisk (*) indicate that the occurrences were counted twice because the same fauna was present across two different stages.

A.

| Region | Tremadocian | Floian | Dapingian | Darriwilian | Sandbian | Katian | Hirnantian |
|---|---|---|---|---|---|---|---|
| Gondwana | 1 | *3 | 2 | *8 | 2 | 2 | 0 |
| Siberia | 0 | 0 | 0 | 1 | 0 | 2 | *3 |
| Laurasia | 0 | 0 | 0 | 1 | 4 | *8 | 0 |
| Baltic | 0 | 0 | 0 | 0 | 0 | 0 | 0 |
| China | 0 | 0 | 0 | 0 | 0 | 0 | 0 |
| Total Sites | 1 | *3 | 2 | *10 | 6 | *12 | *3 |

B.

| Region | Rhuddanian | Aeronian | Telychian | Sheinwoodian | Homerian | Gorstian | Ludfordian | Pridoli |
|---|---|---|---|---|---|---|---|---|
| Gondwana | 0 | 0 | 0 | 0 | 1 | *2 | *2 | 0 |
| Siberia | *4 | *10 | *10 | *9 | *6 | 6 | *7 | *4 |
| Laurasia | 1 | *5 | 7 | *7 | *6 | *12 | *8 | 9 |
| Baltic | 0 | 3 | *8 | 10 | *9 | *11 | *14 | *11 |
| China | 0 | 1 | 11 | *2 | 1 | 1 | 4 | *1 |
| Total Sites | *5 | *19 | *36 | *28 | *22 | *32 | *35 | *25 |

**Table S5. Genus-level richness for jawed and jawless gnathostomes in each region per stage. (A) to (D)** The raw data. **(E) to (H)** Per million years in each stage **(A) and (E)** Ordovician jawless gnathostomes (N = 22). **(B) and (F)** Ordovician jawed gnathostomes (N = 6). **(C) and (G)** Silurian jawless gnathostomes (N = 387). **(D) and (H)** Silurian jawed gnathostomes (N = 97). The values marked with an asterisk (*) indicate that the occurrences counted twice due to the presence of the same fauna across two different stages.

A. (Jawless Gnathostome)

| **Region** | **Tremadocian** | **Floian** | **Dapingian** | **Darriwilian** | **Sandbian** | **Katian** | **Hirnantian** |
|---|---|---|---|---|---|---|---|
| **Gondwana** | 1 | 1 | 1 | 4 | 1 | 2 | 0 |
| **Siberia** | 0 | 0 | 0 | 1 | 0 | 2 | *2 |
| **Laurasia** | 0 | 0 | 0 | 1 | 3 | 3 | 0 |
| **Baltic** | 0 | 0 | 0 | 0 | 0 | 0 | 0 |
| **China** | 0 | 0 | 0 | 0 | 0 | 0 | 0 |

B. (Jawed Gnathostome)

| **Region** | **Tremadocian** | **Floian** | **Dapingian** | **Darriwilian** | **Sandbian** | **Katian** | **Hirnantian** |
|---|---|---|---|---|---|---|---|
| **Gondwana** | 0 | 0 | 0 | 0 | 0 | 0 | 0 |
| **Siberia** | 0 | 0 | 0 | 0 | 0 | 0 | 0 |
| **Laurasia** | 0 | 0 | 0 | 0 | 3 | *3 | 0 |
| **Baltic** | 0 | 0 | 0 | 0 | 0 | 0 | 0 |
| **China** | 0 | 0 | 0 | 0 | 0 | 0 | 0 |

C**.** (Jawless Gnathostome)

| **Region** | **Rhuddanian** | **Aeronian** | **Telychian** | **Sheinwoodian** | **Homerian** | **Gorstian** | **Ludfordian** | **Pridoli** |
|---|---|---|---|---|---|---|---|---|
| **Gondwana** | 0 | 0 | 0 | 0 | 1 | 2 | *4 | 0 |
| **Siberia** | 3 | 4 | *4 | 11 | *10 | 13 | *18 | *15 |
| **Laurasia** | 1 | 2 | 9 | *20 | *15 | 19 | *12 | *27 |
| **Baltic** | 0 | 2 | 3 | 19 | *30 | *37 | *36 | *26 |
| **China** | 0 | 0 | 25 | *8 | 0 | 3 | 5 | *3 |

D. (Jawed Gnathostome)

| **Region** | **Rhuddanian** | **Aeronian** | **Telychian** | **Sheinwoodian** | **Homerian** | **Gorstian** | **Ludfordian** | **Pridoli** |
|---|---|---|---|---|---|---|---|---|
| **Gondwana** | 0 | 0 | 0 | 0 | 0 | 1 | 2 | 0 |
| **Siberia** | 3 | *3 | 6 | *7 | 0 | 0 | 3 | 5 |
| **Laurasia** | 0 | 0 | 1 | 3 | *1 | 0 | 0 | 6 |
| **Baltic** | 0 | 0 | 0 | 0 | 0 | 1 | 5 | *6 |
| **China** | 0 | 2 | 12 | *1 | 1 | 0 | 15 | *14 |

E. (Jawless Gnathostome)

| Region | Tremadocian | Floian | Dapingian | Darriwilian | Sandbian | Katian | Hirnantian |
|---|---|---|---|---|---|---|---|
| **Gondwana** | 0.10 | 0.17 | 0.53 | 0.36 | 0.19 | 0.26 | 0.00 |
| **Siberia** | 0.00 | 0.00 | 0.00 | 0.09 | 0.00 | 0.26 | *0.95 |
| **Laurasia** | 0.00 | 0.00 | 0.00 | 0.09 | 0.56 | 0.39 | 0.00 |
| **Baltic** | 0.00 | 0.00 | 0.00 | 0.00 | 0.00 | 0.00 | 0.00 |
| **China** | 0.00 | 0.00 | 0.00 | 0.00 | 0.00 | 0.00 | 0.00 |

F. (Jawed Gnathostome)

| Region | Tremadocian | Floian | Dapingian | Darriwilian | Sandbian | Katian | Hirnantian |
|---|---|---|---|---|---|---|---|
| **Gondwana** | 0.00 | 0.00 | 0.00 | 0.00 | 0.00 | 0.00 | 0.00 |
| **Siberia** | 0.00 | 0.00 | 0.00 | 0.00 | 0.00 | 0.00 | 0.00 |
| **Laurasia** | 0.00 | 0.00 | 0.00 | 0.00 | 0.56 | *0.39 | 0.00 |
| **Baltic** | 0.00 | 0.00 | 0.00 | 0.00 | 0.00 | 0.00 | 0.00 |
| **China** | 0.00 | 0.00 | 0.00 | 0.00 | 0.00 | 0.00 | 0.00 |

G. (Jawless Gnathostome)

| Region | Rhuddanian | Aeronian | Telychian | Sheinwoodian | Homerian | Gorstian | Ludfordian | Pridoli |
|---|---|---|---|---|---|---|---|---|
| **Gondwana** | 0.00 | 0.00 | 0.00 | 0.00 | 0.26 | 1.18 | *1.74 | 0.00 |
| **Siberia** | 1.15 | 2.11 | *0.70 | 4.78 | *2.56 | 7.65 | *7.83 | *4.84 |
| **Laurasia** | 0.38 | 1.05 | 1.58 | *8.70 | *3.85 | 11.18 | *5.22 | *8.71 |
| **Baltic** | 0.00 | 1.05 | 0.53 | 8.26 | *7.69 | *21.76 | *15.65 | *8.39 |
| **China** | 0.00 | 0.00 | 4.39 | *3.48 | 0.00 | 1.76 | 2.17 | *0.97 |

H. (Jawed Gnathostome)

| Region | Rhuddanian | Aeronian | Telychian | Sheinwoodian | Homerian | Gorstian | Ludfordian | Pridoli |
|---|---|---|---|---|---|---|---|---|
| **Gondwana** | 0.00 | 0.00 | 0.00 | 0.00 | 0.00 | 0.00 | 0.87 | 0.00 |
| **Siberia** | 1.15 | *1.58 | 1.05 | *3.04 | 0.00 | 0.00 | 1.30 | 1.61 |
| **Laurasia** | 0.00 | 0.00 | 0.18 | 1.30 | *0.26 | 0.00 | 0.00 | 1.94 |
| **Baltic** | 0.00 | 0.00 | 0.00 | 0.00 | 0.00 | 0.59 | 2.17 | *1.94 |
| **China** | 0.00 | 1.05 | 2.11 | *0.43 | 0.26 | 0.00 | 6.52 | *4.52 |

**Table S6. Descriptive statistics for faunal composition of assemblages per stage.** Species count, Shannon-Weiner diversity, and taxonomic richness statistics were calculated for all assemblages in each stage**.** Species counts range from 1–40 in 168 assemblages. All site-level values for species count, Shannon diversity, and taxonomic richness are available in Dryad (http://datadryad.org/share/AOhNvKgGWKZHK2LZJ1Q8xi8pl2Z8V_gi4Dk-P4FjTd0).

| Stage | Species Count | | | Shannon-Wienner Count | | | Taxonomic Richness | | |
|---|---|---|---|---|---|---|---|---|---|
| | Mean | SD | Median | Mean | SD | Median | Mean | SD | Median |
| Tremadocian | 1.00 | N/A | 1.00 | 0.00 | N/A | 0.00 | 1.00 | N/A | 1.00 |
| Floian | 1.00 | 0.00 | 1.00 | 0.00 | 0.00 | 0.00 | 1.00 | 0.00 | 1.00 |
| Dapingian | 1.00 | 0.00 | 1.00 | 0.00 | 0.00 | 0.00 | 1.00 | 0.00 | 1.00 |
| Darriwilian | 1.88 | 1.64 | 1.00 | 0.19 | 0.38 | 0.00 | 1.38 | 0.74 | 1.00 |
| Sandbian | 4.17 | 3.87 | 2.50 | 0.50 | 0.59 | 0.32 | 2.00 | 1.26 | 1.50 |
| Katian | 1.33 | 0.71 | 1.00 | 0.07 | 0.21 | 0.00 | 1.11 | 0.33 | 1.00 |
| Hirnantian | 1.00 | 0.00 | 1.00 | 0.00 | 0.00 | 0.00 | 1.00 | 0.00 | 1.00 |
| Rhuddanian | 3.50 | 2.65 | 3.00 | 0.46 | 0.61 | 0.28 | 2.00 | 1.41 | 1.50 |
| Aeronian | 3.00 | 2.39 | 2.00 | 0.00 | 0.00 | 0.00 | 1.00 | 0.00 | 1.00 |
| Telychian | 5.45 | 4.29 | 5.00 | 0.27 | 0.37 | 0.00 | 1.52 | 0.74 | 1.00 |
| Sheinwoodian | 9.00 | 8.04 | 7.00 | 0.47 | 0.43 | 0.56 | 2.09 | 1.08 | 2.00 |
| Homerian | 5.13 | 5.91 | 3.00 | 0.23 | 0.43 | 0.00 | 1.50 | 0.93 | 1.00 |
| Gorstian | 10.74 | 10.90 | 7.00 | 0.28 | 0.36 | 0.00 | 1.78 | 1.15 | 1.00 |
| Ludfordian | 10.65 | 11.00 | 8.00 | 0.53 | 0.60 | 0.00 | 2.35 | 1.58 | 1.00 |
| Pridoli | 10.13 | 10.36 | 5.50 | 0.52 | 0.52 | 0.40 | 2.31 | 1.35 | 2.00 |

**Table S7.**
**CCA results**. Results table be found in the Dryad folder (http://datadryad.org/share/AOhNvKgGWKZHK2LZJ1Q8xi8pl2Z8V_gi4Dk-P4FjTd0.). Eigenvalues for the correspondence axes are as follows: **axis 1 = 0.39**, **axis 2 = 2.65 × $10^{-2}$** **(**permutation N = 1,000,000). For site scores, refer to Fig. 2B **Left** for the plot. For taxon scores, refer to Fig. 2B **Right** for the plot.

**Table S8.**
**Factor analysis of all sites.** Results table and details of the dataset used for this analysis are available on Dryad at (http://datadryad.org/share/AOhNvKgGWKZHK2LZJ1Q8xi8pl2Z8V_gi4Dk-P4FjTd0.). Site Score and Taxon loading for Fig. S9 plot.

**Table S9.**
**ANOSIM results for comparison of assemblages across geological series boundaries** (permutation N = 1,000,000). **(A)** R statistics for raw species richness within groups using different dissimilarity metrics. Values closer to 0 indicate greater similarity between faunas, while values closer to 1 indicate high dissimilarity. **(B)** p-values. Significant values are marked with an asterisk (*) (α = 0.05).

A.

| Comparison | Over all Mean | Abundance Mean | Bray-Curtis | Kulczynski | Chord | Morisita | Raup-Crick |
|---|---|---|---|---|---|---|---|
| Middle Orddovician - Upper Ordovician | 0.52 | 0.62 | 0.52 | 0.64 | 0.46 | 0.39 | 0.59 |
| Upper Ordovician - Llandovery Silurian | 0.38 | 0.45 | 0.39 | 0.39 | 0.31 | 0.28 | 0.50 |
| Llandovery Silurian - Wenlock Silurian | 0.14 | 0.17 | 0.14 | 0.12 | 0.11 | 0.09 | 0.22 |
| Wenlock Silurian -Ludlow Silurian | -0.03 | -0.02 | -0.03 | -0.03 | -0.03 | -0.05 | -0.02 |
| Ludlow Silurian - Pridoli Silurian | 0.08 | 0.09 | 0.07 | 0.08 | 0.09 | 0.07 | 0.10 |

B.

| Comparison | Over all Mean | Abundance Mean | Bray-Curtis | Kulczynski | Chord | Morisita | Raup-Crick |
|---|---|---|---|---|---|---|---|
| Middle Orddovician - Upper Ordovician | 0.11 | 0.07 | 0.13 | 0.07 | 0.13 | 0.13 | 0.07 |
| Upper Ordovician - Llandovery Silurian | 0.02 | 0.01 | 0.01 | 0.01 | 0.02 | 0.03 | <0.01* |
| Llandovery Silurian - Wenlock Silurian | 0.01 | <0.01* | <0.01* | 0.01 | 0.01 | 0.02 | <0.01* |
| Wenlock Silurian -Ludlow Silurian | 0.81 | 0.72 | 0.83 | 0.81 | 0.84 | 0.93 | 0.63 |
| Ludlow Silurian - Pridoli Silurian | 0.15 | 0.13 | 0.17 | 0.16 | 0.14 | 0.18 | 0.10 |

**Table S10.**
**SIMPER total average dissimilarities.** Results based on raw diversity data.

| **Boundary** | **Middle Ordovician - Upper Ordovician** | **Upper Ordovician - Llandovery Silurian** | **Llandovery Silurian - Wenlock Silurian** | **Wenlock Silurian - Ludlow Silurian** | **Ludlow Silurian - Pridoli Silurian** |
|---|---|---|---|---|---|
| **Average Dissimilarity** | 77.66 | 90.58 | 66.04 | 62.07 | 71.22 |

**Table S11. SIMPER taxon loadings based on raw diversity data in assemblages across each geological series boundary. (A)** Middle Ordovician and Upper Ordovician. **(B)** Upper Ordovician and Llandovery Silurian. **(C)** Llandovery Silurian and Wenlock Silurian. **(D)** Wenlock Silurian and Ludlow Silurian. **(E)** Ludlow Silurian and Pridoli.

A.

| Taxon | Av. dissim | Contrib. % | Cumulative % | Mean abound.Mid | Mean abound.Upp |
|---|---|---|---|---|---|
| Arandaspida | 25.19 | 32.43 | 32.43 | 2.50 | 0.00 |
| Astraspida | 19.08 | 24.56 | 56.99 | 0.50 | 2.50 |
| Unknown | 14.07 | 18.11 | 75.10 | 1.50 | 1.25 |
| Pteraspidomorpha | 7.36 | 9.47 | 84.57 | 0.00 | 0.75 |
| Chondrichthyes | 6.91 | 8.89 | 93.47 | 0.00 | 1.00 |
| Thelodonti | 5.07 | 6.53 | 100.00 | 0.00 | 0.50 |
| Osteichthyes | 0.00 | 0.00 | 100.00 | 0.00 | 0.00 |
| Anaspida | 0.00 | 0.00 | 100.00 | 0.00 | 0.00 |
| Placoderm | 0.00 | 0.00 | 100.00 | 0.00 | 0.00 |
| Acanthodii | 0.00 | 0.00 | 100.00 | 0.00 | 0.00 |
| Galeaspidomorphi | 0.00 | 0.00 | 100.00 | 0.00 | 0.00 |
| Osteostraci | 0.00 | 0.00 | 100.00 | 0.00 | 0.00 |
| Heterostraci | 0.00 | 0.00 | 100.00 | 0.00 | 0.00 |

B.

| Taxon | Av. dissim | Contrib. % | Cumulative % | Mean abound.Upp | Mean abound.Lla |
|---|---|---|---|---|---|
| Thelodonti | 28.25 | 31.18 | 31.18 | 0.50 | 3.34 |
| Astraspida | 21.16 | 23.36 | 54.54 | 2.50 | 0.00 |
| Chondrichthyes | 13.92 | 15.36 | 69.91 | 1.00 | 1.66 |
| Galeaspidomorphi | 9.63 | 10.63 | 80.53 | 0.00 | 1.28 |
| Unknown | 8.46 | 9.34 | 89.87 | 1.25 | 0.03 |
| Pteraspidomorpha | 6.34 | 7.00 | 96.88 | 0.75 | 0.00 |
| Acanthodii | 1.59 | 1.75 | 98.63 | 0.00 | 0.21 |
| Heterostraci | 0.78 | 0.86 | 99.48 | 0.00 | 0.10 |
| Anaspida | 0.26 | 0.29 | 99.77 | 0.00 | 0.03 |
| Placodermi | 0.21 | 0.23 | 100.00 | 0.00 | 0.03 |
| Osteichthyes | 0.00 | 0.00 | 100.00 | 0.00 | 0.00 |
| Osteostraci | 0.00 | 0.00 | 100.00 | 0.00 | 0.00 |
| Arandaspida | 0.00 | 0.00 | 100.00 | 0.00 | 0.00 |

C.

| Taxon | Av. dissim | Contrib. % | Cumulative % | Mean abound.Lla | Mean abound.Wen |
|---|---|---|---|---|---|
| **Thelodonti** | 28.44 | 43.06 | 43.06 | 3.34 | 7.05 |
| **Anaspida** | 9.62 | 14.57 | 57.62 | 0.03 | 1.95 |
| **Chondrichthyes** | 9.37 | 14.19 | 71.81 | 1.66 | 0.32 |
| **Galeaspidomorphi** | 7.66 | 11.59 | 83.41 | 1.28 | 0.00 |
| **Heterostraci** | 4.94 | 7.48 | 90.89 | 0.10 | 0.59 |
| **Osteostraci** | 4.13 | 6.25 | 97.14 | 0.00 | 0.86 |
| **Acanthodii** | 1.49 | 2.26 | 99.40 | 0.21 | 0.05 |
| **Unknown** | 0.22 | 0.34 | 99.74 | 0.03 | 0.00 |
| **Placodermi** | 0.17 | 0.26 | 100.00 | 0.03 | 0.00 |
| **Osteichthyes** | 0.00 | 0.00 | 100.00 | 0.00 | 0.00 |
| **Astraspida** | 0.00 | 0.00 | 100.00 | 0.00 | 0.00 |
| **Arandaspida** | 0.00 | 0.00 | 100.00 | 0.00 | 0.00 |
| **Pteraspidomorpha** | 0.00 | 0.00 | 100.00 | 0.00 | 0.00 |

D.

| Taxon | Av. dissim | Contrib. % | Cumulative % | Mean abound.Wen | Mean abound.Lud |
|---|---|---|---|---|---|
| **Thelodonti** | 31.70 | 51.07 | 51.07 | 7.05 | 9.28 |
| **Anaspida** | 9.52 | 15.33 | 66.40 | 1.95 | 1.50 |
| **Osteostraci** | 6.60 | 10.63 | 77.02 | 0.86 | 1.22 |
| **Heterostraci** | 6.27 | 10.11 | 87.13 | 0.59 | 0.81 |
| **Acanthodii** | 3.35 | 5.40 | 92.53 | 0.05 | 0.69 |
| **Galeaspidomorphi** | 1.79 | 2.89 | 95.42 | 0.00 | 0.25 |
| **Chondrichthyes** | 1.51 | 2.43 | 97.84 | 0.32 | 0.03 |
| **Osteichthyes** | 0.86 | 1.38 | 99.23 | 0.00 | 0.34 |
| **Placodermi** | 0.48 | 0.77 | 100.00 | 0.00 | 0.13 |
| **Unknown** | 0.00 | 0.00 | 100.00 | 0.00 | 0.00 |
| **Astraspida** | 0.00 | 0.00 | 100.00 | 0.00 | 0.00 |
| **Arandaspida** | 0.00 | 0.00 | 100.00 | 0.00 | 0.00 |
| **Pteraspidomorpha** | 0.00 | 0.00 | 100.00 | 0.00 | 0.00 |

E.

| Taxon | Av. dissim | Contrib. % | Cumulative % | Mean abound.Lud | Mean abound.Pri |
|---|---|---|---|---|---|
| **Thelodonti** | 33.98 | 47.70 | 47.70 | 9.28 | 7.00 |
| **Heterostraci** | 12.98 | 18.22 | 65.93 | 0.81 | 2.58 |
| **Acanthodii** | 7.46 | 10.47 | 76.40 | 0.69 | 1.92 |
| **Osteostraci** | 6.38 | 8.96 | 85.36 | 1.22 | 0.75 |
| **Anaspida** | 6.33 | 8.89 | 94.25 | 1.50 | 0.42 |
| **Galeaspidomorphi** | 1.67 | 2.34 | 96.59 | 0.25 | 0.00 |
| **Osteichthyes** | 1.54 | 2.16 | 98.74 | 0.34 | 0.17 |
| **Placodermi** | 0.45 | 0.64 | 99.38 | 0.13 | 0.00 |
| **Chondrichthyes** | 0.44 | 0.62 | 100.00 | 0.03 | 0.08 |
| **Unknown** | 0.00 | 0.00 | 100.00 | 0.00 | 0.00 |
| **Astraspida** | 0.00 | 0.00 | 100.00 | 0.00 | 0.00 |
| **Arandaspida** | 0.00 | 0.00 | 100.00 | 0.00 | 0.00 |
| **Pteraspidomorpha** | 0.00 | 0.00 | 100.00 | 0.00 | 0.00 |

**Table S12. Canonical Correspondence Analysis (CCA) for alternative assignments of Chondrichthyes and Acanthodi (Reanalysis 1–3)**. Results table and details of the dataset used for this analysis are available on Dryad at (http://datadryad.org/share/AOhNvKgGWKZHK2LZJ1Q8xi8pl2Z8V_gi4Dk-P4FjTd0.). **Reanalysis 1:** Eigenvalues for the correspondence axes are as follows: **axis 1 = 0.39**, **axis 2 = $2.39 \times 10^{-2}$** (permutation N = 1,000,000). For site scores, refer to Fig. S9A left for the plot; for taxon scores, refer to Fig. S9A right for the plot. **Reanalysis 2:** Eigenvalues for the correspondence axes are as follows: **axis 1 = 0.29**, **axis 2 = $2.62 \times 10^{-2}$** (permutation N = 1,000,000). For site scores, refer to Fig. S11 left for the plot; for taxon scores, refer to Fig. S11 right for the plot. **Reanalysis 3:** Eigenvalues for the correspondence axes are as follows: **axis 1 = 0.30**, **axis 2 = $2.39 \times 10^{-2}$** (permutation N = 1,000,000). For site scores, refer to Fig. S14 left for the plot; for taxon scores, refer to Fig. S14 right for the plot.

**Table S13.**
**Factor Analysis for Ordovician Chondrichthyes reassigned as "Unknown" (reanalysis 1)**
Results table and details of the dataset used for this analysis are available on Dryad at (http://datadryad.org/share/AOhNvKgGWKZHK2LZJ1Q8xi8pl2Z8V_gi4Dk-P4FjTd0.). This table presents a reanalysis of factor analysis where all Ordovician Chondrichthyes are categorized as Unknown. Site Score and Taxon loading for Fig. S9E–G plot.

**Data S1. (separate file)**

**Assemblage descriptions and gnathostome faunal lists.** Detailed all faunal information from Ordovician to Silurian (N = 168) and reference literatures.

**Data S2. (separate file)**

**Matrix of gnathostome occurrence in all assemblages.** Input matrix for histograms and Tab. S4.

**Data S3. (separate file)**

**Ordovician gnathostome occurrences.** Raw occurrence records from the reference literature for the Ordovician.

**Data S4. (separate file)**

**Silurian gnathostome occurrence.** Raw occurrence records from the the reference literature for Silurian.

**Data S5. (separate file)**

**Devonian gnathostome occurrence.** Raw occurrence records from the reference literature for Devonian. These were not included in our assemblage analyses.

**Data S6. (separate file)**

**Ordovician conodont occurrences.** Raw occurrence records from "The Paleobiology Database" website (*34*) for each stage in the Ordovician.

**Data S7. (separate file)**

**Silurian conodont occurrence.** Raw occurrence records from "The Paleobiology Database" website (*34*) for each stage in the Silurian.

**Data S8. (separate file)**

**Matrix of gnathostome occurrences for statistical analysis.** Input matrix for statistical analysis including cluster analysis, CCA, NMDS, FA, ANOSIM and SIMPER. Assemblages with less than 3 species excluded.
